# Cross-Modal Representation of Identity in Primate Hippocampus

**DOI:** 10.1101/2022.09.12.507611

**Authors:** Timothy J Tyree, Michael Metke, Cory T Miller

## Abstract

Faces and voices are the dominant social signals used to recognize individuals amongst human and nonhuman primates (*1*–*4*). Yet it is not known how these critical signals are integrated into a cross-modal representation of individual identity in the primate brain. Here we show that while, like humans (*5*–*7*), single neurons in the marmoset hippocampus exhibit selective responses when presented with the face or voice of a specific individual conspecific, a parallel mechanism for representing the cross-modal identities for multiple individuals is evident within single neurons and at a population level. Manifold projections likewise showed separability of individuals, as well as clustering for others’ families, suggesting that multiple learned social categories are encoded as related dimensions of identity in hippocampus. These findings demonstrate that neural representations of identity in hippocampus are both modality-independent and reflect the hierarchical structure of the primate social network.

**One-Sentence Summary:** We show that cross-modal representations of identity in primate hippocampus can be achieved by at least two distinct neural mechanisms and comprise multiple social categories that reflect different relationships.

## Main Text

Navigating the complex societies that typify primates relies learning the identity of each individual in the group and their respective social relationships through observation (*8*). Although evidence shows that neurons in the brains of primates and other mammals selectively respond to the identity when viewing the face or hearing the voice of a specific individual as unimodal signals (*2, 9*–*13*), data showing that single neurons are responsive to both the face and voice of an individual – a cross-modal representation of identity – is limited to ‘concept cells’ in human hippocampus; a sparse population of highly-selective neurons responsive to well-known individuals and locations across different views and modalities learned through observation (*5*–*7*). These neurons are significant for several reasons including their putative role in memory functions (*14*) and potential uniqueness to humans (*15*). Here we tested whether cross-modal representations of identity are evident in the hippocampus of marmoset monkeys by recording single hippocampal neurons (*16*) while presenting subjects with multiple exemplars of individual marmoset faces - from different viewpoints - and voices as unimodal stimuli, consistent with previous work (*10*) as well as concurrently by presenting the faces and voices from the same (identity match) or different individuals (identity mismatch): i.e. Match versus Mismatch (MvMM). Visual stimuli were presented from a monitor directly in front of the animal while a speaker positioned directly below the screen broadcasted the acoustic stimuli. Subjects were only presented with familiar conspecifics housed in the same colony that differed in their respective social relatedness (e.g. family members and non-family members (*7*)).

To first test whether cross-modal representations of identity are evident in the hippocampus of a nonhuman primate, we performed the same ROC selectivity analysis described previously in humans (*5*– *7*) and revealed a population of cross-modal invariant neurons for individual identity when observing marmoset faces or voices (Figure 1A, S1A), as well as neurons selective for individual identity when viewing only their faces (Figure 1B, S1B), or hearing only their voices (Figure 1C, S1C). These identity neurons were confirmed in all hippocampal subfields (Figure 1D). Overall, we observed N=148 (9.2%) of N=1,602 qualifying neurons demonstrated selectivity for a single preferred individual (Figure 1E) with different neurons selective for faces (N=52), voices (N=39) or both faces and voices (N=57; Figure 1F). The mean area under the ROC curve (AUC) of identity neurons (AUC=0.902±0.014) was significantly above chance (p<0.001, Figure 1G). Although these neurons in marmosets were overall less selective than in humans (*5*–*7*), this disparity may reflect species differences in the baseline hippocampus activity (Figure S2) that affect neural coding mechanisms for identity.

**Figure 1.**
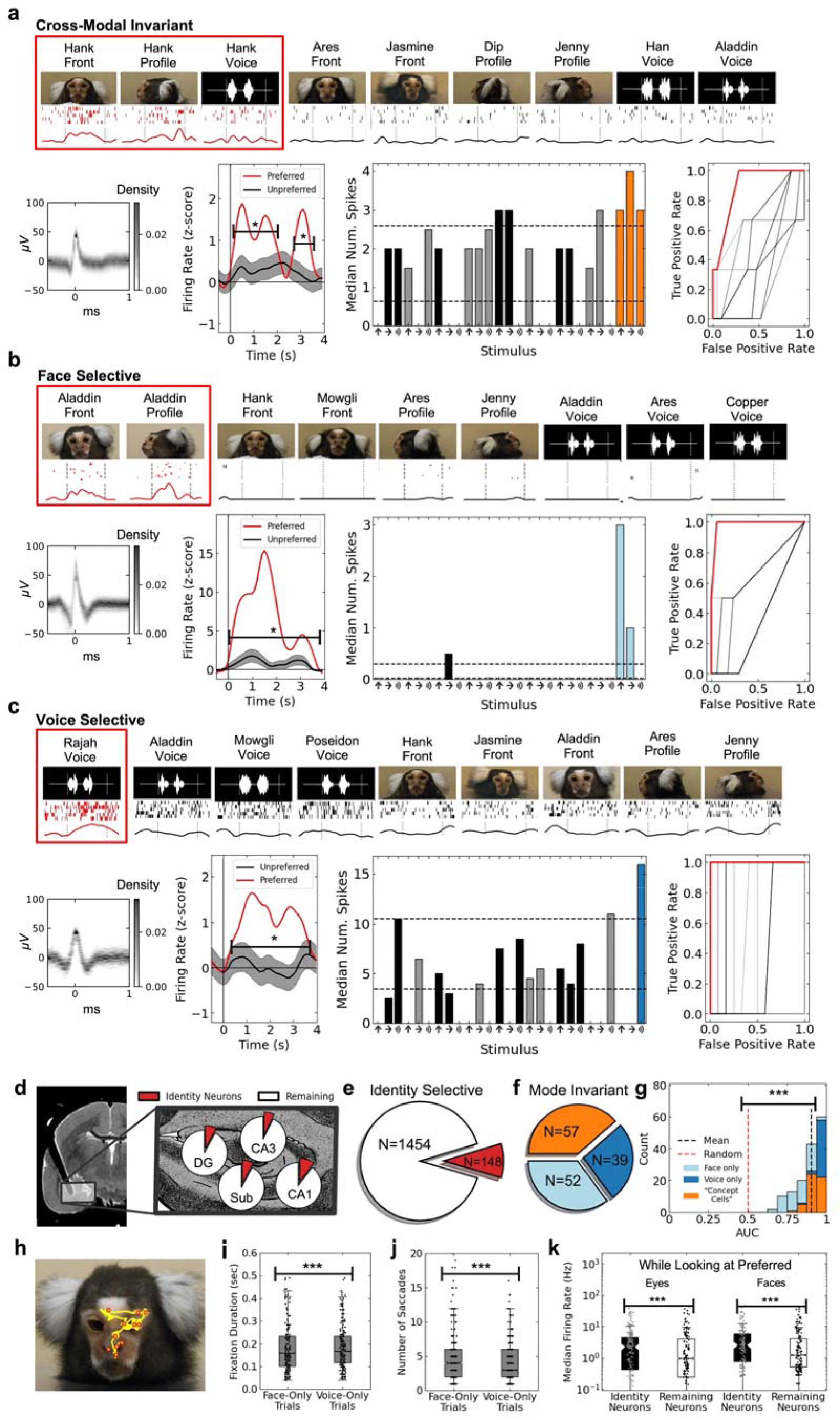
Putative ‘concept cells’ in marmoset hippocampus. **[a-c]** Top row: subset of stimuli shown above raster and PSTH. Bottom row: spike waveform density; normalized PSTH to all stimuli (preferred: red, nonpreferred: black), indicated are time points that show significant difference (p<0.05); median number of spikes for unimodal stimuli (grey/black indicate non-preferred individuals; ROC curve (shuffled controls shown in black). **E**xemplar identity neurons responding selectively to [a] the face and voice of a preferred conspecific (red), [b] the face only, and [c] the voice only. **[d]** Anatomical distribution of identity neurons (red) in hippocampal subfields relative to neurons remaining that responded to any stimulus (white). Black shadow indicates the electrode array track with MRI distortion artifact. **[e]** Pie chart showing the abundance of identity neurons in black with the number of remaining neurons that qualified for the ROC selectivity analysis in white. **[f]** Pie chart showing the mode distribution of identity neurons. Modes included face (light blue), voice (dark blue), and both (orange). **[g]** Histogram showing the distribution of areas under ROC curves comparable to red ROC curves in [a-c]. Colors are as in [f]. Black dotted line is the mean, while red dotted line is the mean of 10,000 random shuffles of the labels. **[h]** Exemplar eye-movements (yellow) with fixations indicated (red). **[i]** Distribution of eye fixation durations for unimodal trials. **[j]** Distribution of apparent saccade number for unimodal trials. **[k]** Distribution of median firing rates while observer was looking at eyes (left) and face (right) for identity neurons (black) *versus* remaining neurons (white). Three asterisks indicate a significant median difference (p<0.001).

Analysis of eye-movements (Figure 1H) revealed marmosets’ visual behavior and neural activity were differentially affected by modality and identity. Marmosets exhibited significantly shorter fixations (p<0.001, N_fixations_=18,965; Figure 1I) and significantly more saccades (p<0.001, N_saccades_=2,203) during trials with face-only relative to the voice-only trials (Figure 1J). These monkeys were also highly focused on faces during stimulus presentations, with faces accounting for 77.9% of viewing time and eyes specifically accounted for 37.6% of viewing time. The firing rate of identity neurons was significantly greater than the remaining neurons when subjects were looking at the eyes or face (both p<0.001; Figure 1K) suggesting that this class of neurons was particularly sensitive to faces and facial features regardless of identity. This was not, however, a broad attentional effect (*17*), as the firing rate of simultaneously recorded non-identity neurons did not show the same increased firing rate when gazing at faces or eyes.

A potential parallel mechanism to highly-selective “concept cells” is for individual cells to contribute to multiple functions (*18, 19*), such as single neurons being sensitive to the cross-modal identity of multiple conspecifics. Previous studies show that hippocampal neurons are sensitive to mismatches between the features of a particular stimulus and previously learned category (*20, 21*). To test whether a similar mechanism is evident for the learned social identities of conspecifics in marmoset hippocampus, we next analyzed whether neurons would respond differently when simultaneously observing the face and voice from the same (identity match) or different (identity mismatch) individuals. By presenting a face and voice in all MvMM trials, we controlled for the potential effects of multi-modal integration (Figure S3A) and instead tested whether a subordinate category– identity– elicited changes in neural activity. Analyses revealed that indeed a subpopulation of units– MvMM neurons– exhibited a significant firing rate preference for either match trials (Figure 2A) or mismatch trials (Figure 2B), with some neurons modulated only by this category distinction (Figure 2A) and others more generally stimulus drive (Figure 2B). Overall, 21.7% of neurons (N=511 of 2,358) exhibited a significant response during MvMM trials, with significantly more units exhibiting a higher firing rate during match (N=401) than mismatch (N=110) trials (p<0.001; Figures 2C, S3B). MvMM neurons were largely distinct from the identity neurons described above (Figure 2D). Interestingly, 56% of the neurons observed in both populations whose anatomical location could be confirmed were recorded in CA1. In contrast to identity neurons, MvMM neurons were biased to CA1 (Figure 2E), with N=155 (44.3%) out of 350 neurons confirmed in the CA1 qualifying as MvMM neurons. In CA1, significantly more MvMM neurons (N=129/155, 83.2%) preferred match trials to mismatch trials (p<0.001). Analysis of visual behavior revealed that the MvMM neurons exhibited significantly higher median firing rate while the subject was looking at the eyes or face (p<0.001, N=511; Figure 2F) indicating that these neurons were likewise sensitive to these socially-relevant features. Further analysis indicated that marmosets exhibited significantly more saccadic eye movements during mismatch trials (Figure 2G) and that this difference in behavior was most prominent 1-2s after stimulus onset (p<0.05, N_saccades_=4,603; Figure 2H) suggesting that the monkeys were perceptually sensitive to the incongruence in the subordinate category– identity– shared between cross-modal signals, consistent with the pattern of neural responses to these stimuli.

**Figure 2.**
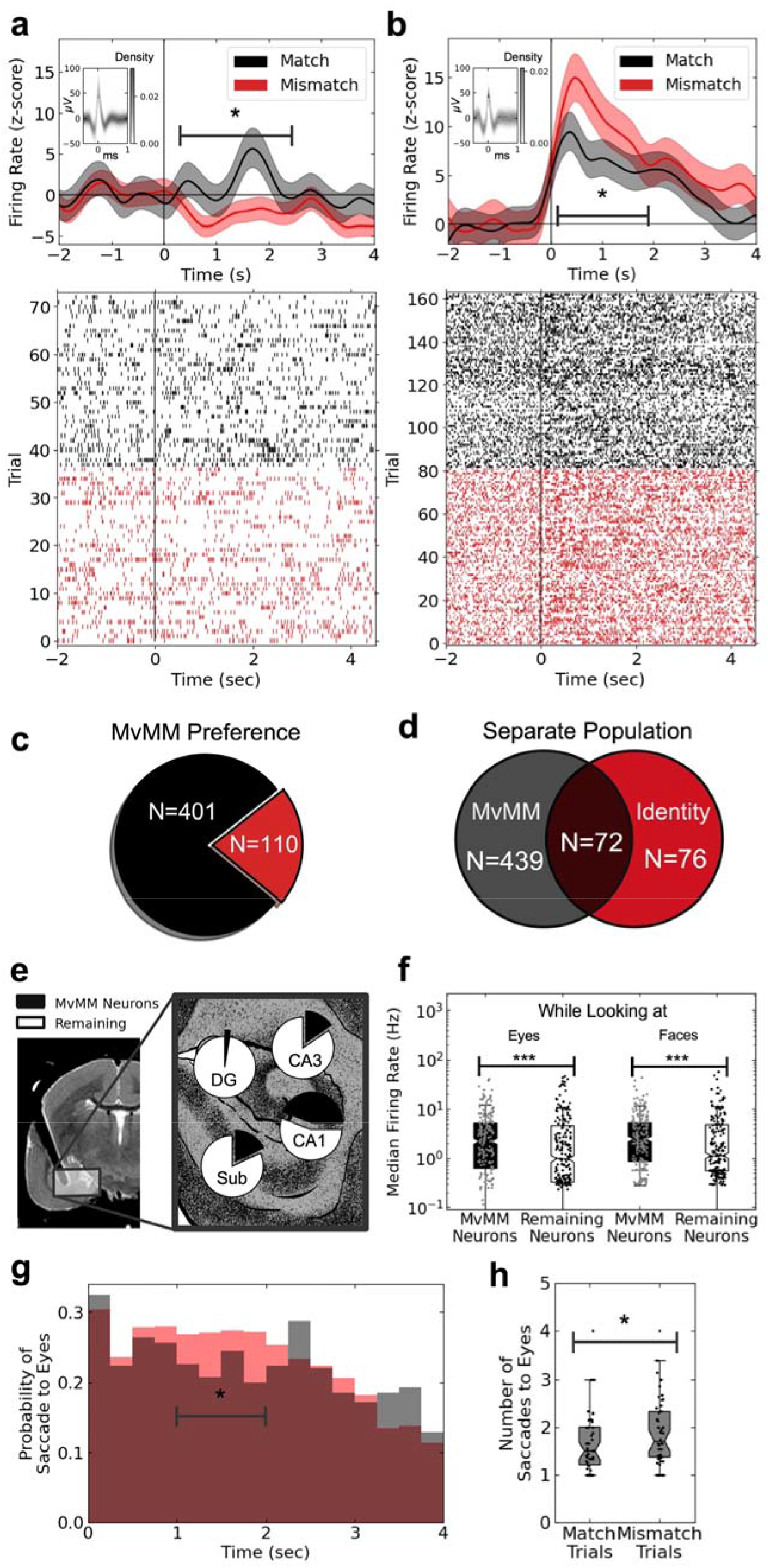
Single neurons in hippocampus represent multiple individuals. **[a**,**b]** The PSTH normalized by the pre-stimulus baseline (top) and spike raster (bottom) for two exemplar MvMM neurons. Black indicates match and red indicates mismatch trials. Vertical line indicates stimulus onset. Inset shows spike waveform density. Asterisk indicates significant time points (p<0.05). Exemplar neuron with higher firing rate for [a] match and [b] mismatch trials. **[c]** Pie chart showing the number of neurons that responded significantly more for match (black) or mismatch (red) trials. **[d]** Venn diagram showing the number of MvMM neurons (black) in common with identity neurons (red). **[e]** Relative abundance of MvMM neurons in each hippocampal subfield. **[f]** Distribution of median firing rate while looking at the eyes (left) and face (right) for MvMM neurons (black) *versus* remaining neurons (white). **[g]** Probability density of saccadic eye movements directed towards the eyes for match (black) and mismatch (red) trials. Indicated are the time points in [h]. **[h]** Distribution of apparent number of saccades to eyes. Asterisk indicates significant median difference (p<0.05).

These findings suggest two seemingly distinct mechanisms for representing cross-modal identity are evident in primate hippocampus. We conjectured that more temporally selective coding mechanisms in hippocampus may inform how these two processes for encoding identity are integrated at a population level. To test this, we developed an algorithm to identify intervals of time during which individual neurons exhibited significant differences in median firing rate for a specific category (p<0.05), which we labeled as predictive time bins (Figure S4). Importantly, this algorithm was applied to all neurons in the population, not only those classified as identity selective or MvMM neurons (e.g. Figures 1&2). We first implemented this analysis to test whether predictive time bins were selective for specific individuals when observing their face or voice. Figure 3A shows a pair of exemplar neurons that exhibited separate predictive time bins for two different individuals. Analyses revealed that N=1,634 out of 2,358 hippocampal neurons (69.3%) exhibited at least one identity-specific predictive time bin, with the majority comprising predictive time bins for two or more individuals (Figure 3B). Identity-specific predictive time bins exhibited a mean AUC (AUC=0.802±0.003) that was significantly above chance (p<0.001, N_bins_=3,958; Figure 3C). Analysis of visual behavior showed that neurons possessing identity-specific predictive time bins exhibited a significant increase in median firing rate when subjects were looking at the face of the preferred individual (Figure 3D). Notably, instances of face and eye viewing were highly variable and not limited to the timing of predictive time bins suggesting that attentional effects from visual behavior were not likely driving neural activity during these periods (Figure S5). We applied the same algorithm to test for predictive time bins that distinguished MvMM trials and found a similar result (Figure 3E) with 1,455 neurons exhibiting MvMM predictive time bins. Furthermore, the firing rate of neurons with predictive time bins for MvMM exhibited a significantly higher firing rate than other neurons when subjects looked at the face than the remaining neurons (p<0.001; Figure 3F). We observed considerable overlap between neurons with identity-specific and MvMM predictive time bins, as 82.2 % (N=1,196, Figure S6A) exhibited predictive time bins in both analyses. These results demonstrate that information about specific identities is evident in the activity of hippocampal neurons using this more temporally refined predictive-time bin analysis.

**Figure 3.**
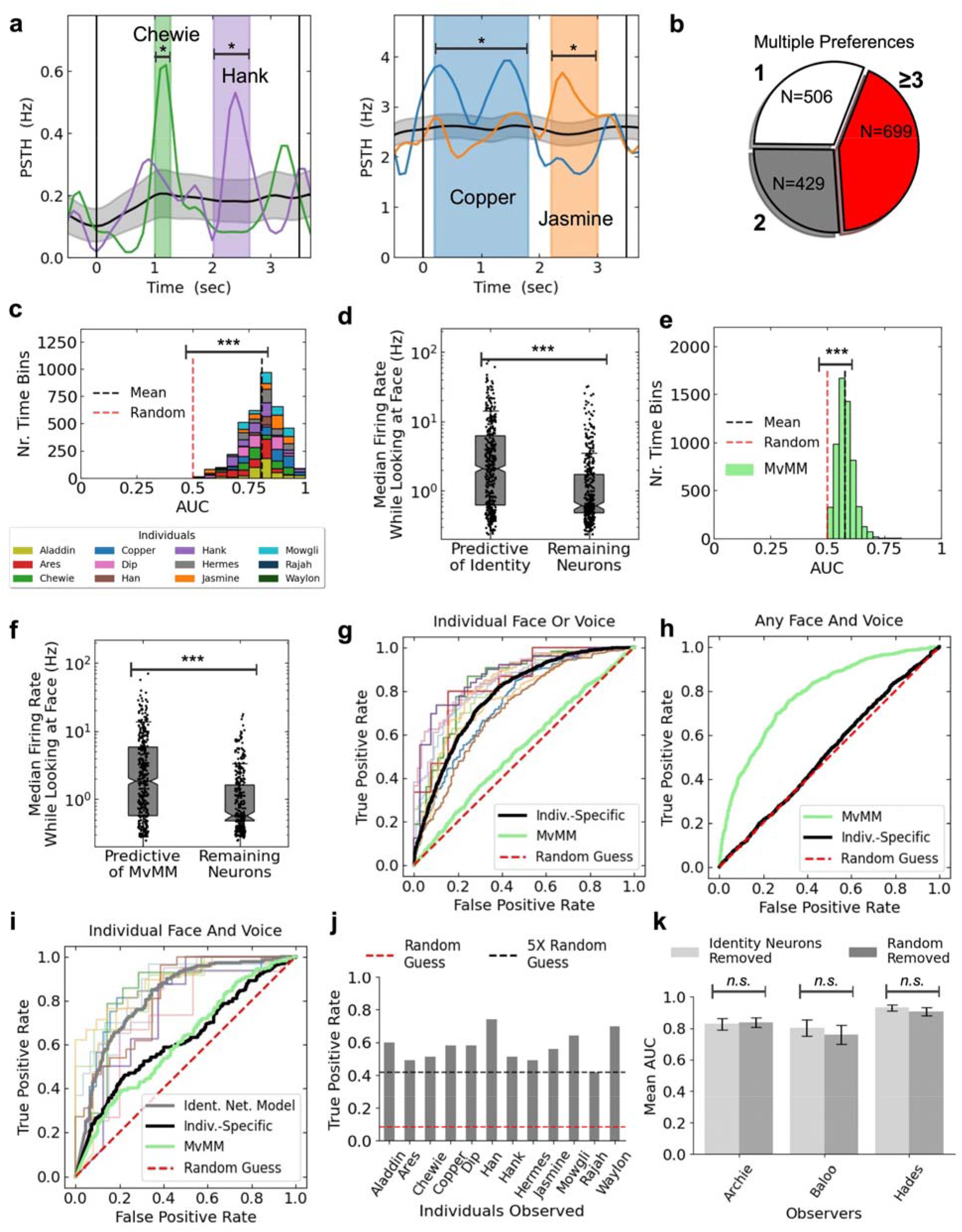
Cross-modal decoding of identity. **[a]** PSTH of two exemplar predictive neurons. Colored traces average over trials involving preferred individual while the gray shaded regions indicate 95% confidence intervals of the session mean. Colored regions indicate identity-specific time bins. **[b]** Pie chart showing number of identity-specific predictive neurons that prefer one individual (white), two individuals (gray), and three or more individuals (red). **[c]** Histogram showing AUC distribution of identity-specific time bins with colors indicating preferred individuals in legend. Dotted lines indicate the mean (black) and the control with shuffled labels (red). **[d]** Distribution of median firing rates while the observer was looking at the face for the identity-specific predictive neurons compared to the remaining neurons. **[e]** Histogram showing AUC distribution of MvMM time bins. Dashed lines indicate the mean (black) and the control with shuffled labels (red). **[f]** Distribution of median firing rates while the observer was looking at the face for the MvMM predictive neurons compared to the remaining neurons. **[g]** ROC curves for the detection of face or voice of individuals. Firing rates were considered from MvMM time bins (green, AUC=0.536) and identity-specific time bins (black, AUC=0.779) similarly averaged over individuals. Thinner colored lines indicate individuals as in [c]. **[h]** ROC curves for the detection of match trials. Firing rates were considered from MvMM time bins (green, AUC=0.782) and from identity-specific time bins (black, AUC=0.516). **[i]** ROC curves for the detection of both face and voice of individuals from same 19 recording sessions as in [g,h]. Firing rates were considered from MvMM time bins (green, AUC=0.615), identity-specific time bins (black, AUC=0.622), and the INM (gray, AUC=0.818), similarly averaged over individuals. Results of the INM for individuals are shown by thin lines colored as in the legend of [c]. Red dotted line indicates random guess as in [g,h]. **[j]** Bar plot showing true positive rates predicted by a winner-take-all model that considered predictions from the INM specific to twelve individuals. Indicated is the mean of the shuffled labels (red) and 5× that value (black). Bar plots summarize the trials from the testing sets of 33 recording sessions (N_trials_=454). **[k]** Bar plot showing mean AUC with identity neurons removed (light gray) *versus* the control randomly removing an equal number of bins from the remaining cells (dark gray). Uncertainty indicates 95% confidence of the mean. No significant difference was observed across recording sessions for any of the three qualifying subjects according to a paired Wilcoxon-Mann-Whitney test (Archie, p=0.81, N_identities_=14; Baloo, p=0.58, N_identities_=9; Hades, p=0.50, N_identities_=12). Three asterisks indicate statistical significance (p<0.001).

Encouraged by these findings, we developed a stable neural decoder by combining the firing rates of predictive time bins using an ensemble of gradient-boosted decisions trees (*22*). When using identity-specific time bins, we could reliably decode the identity of all marmosets when subjects observed their face or voice (accuracy: 77.4%; Figure 3G). Likewise, the same approach could successfully decode MvMM trials when using MvMM time bins (accuracy: 75.7%; Figure 3H). Interestingly, the two kinds of decoders used mostly different time points, with only 24.6%±1.5% of identity-specific time bins overlapping with MvMM time bins within the same neurons (Figure S6B).

To test whether the same population could represent multiple cross-modal identities, we developed the identity network model (INM) that integrates these two decoding approaches. The first approach was identical to the identity-specific decoder described above, resulting in accurate decoding for each individual’s face or voice. The second approach classified MvMM trials as either match or mismatch but was blind to individual identity. Our INM combined these two approaches to achieve cross-modal decoding of individual identity (Figure S7). This combination was critical because the identity-specific predictive population was only accurate for individual identity but performed poorly for classifying MvMM (Figure 3G), while the MvMM predictive population was the inverse (Figure 3H). When combined across individuals, the INM successfully decoded the cross-modal identity of all twelve individuals (accuracy: 84.5%; Figure 3I). Notably, decoding performance was at least 5× above chance when distinguishing all individuals (Figures 3J, S8). Together, these results demonstrate cross-modal representations for the individual identities of multiple conspecifics are evident at the population-level in primate hippocampus (*23*).

Because identity neurons were included in decoding, we investigated whether their explanatory contribution was disproportionate to their sparse distribution. We compared INM performance when these neurons were removed from the analysis and separately used only in the analysis *versus* an equal number of other neurons, and we observed no significant effect on decoding performance despite the consideration of only individuals preferred by identity neurons (Figure 3K, S9-10) suggesting that these highly-selective neurons are no more significant for decoding the cross-modal identity of familiar individuals than other neurons in the population.

The success of the INM provided compelling evidence that an individual within a marmoset’s social network can be decoded from their face and/or voice, but an individual’s identity is also coupled to their social relationships, such as their family. To test whether hippocampus encodes categorical attributes of social identity, we applied nonlinear dimensionality reduction techniques shown to be powerful tools for revealing elements of brain functions (*24*), including in studies of hippocampus (*25*). As a first step to this end, we replicated the findings of the INM using the same identity-specific predictive time bins for marmoset faces and voices drawn from the entire hippocampal population and showed that manifold projections similarly separated individuals (Figure 4A, S11), including for different subpopulations of neurons (Figure S11G).

**Figure 4.**
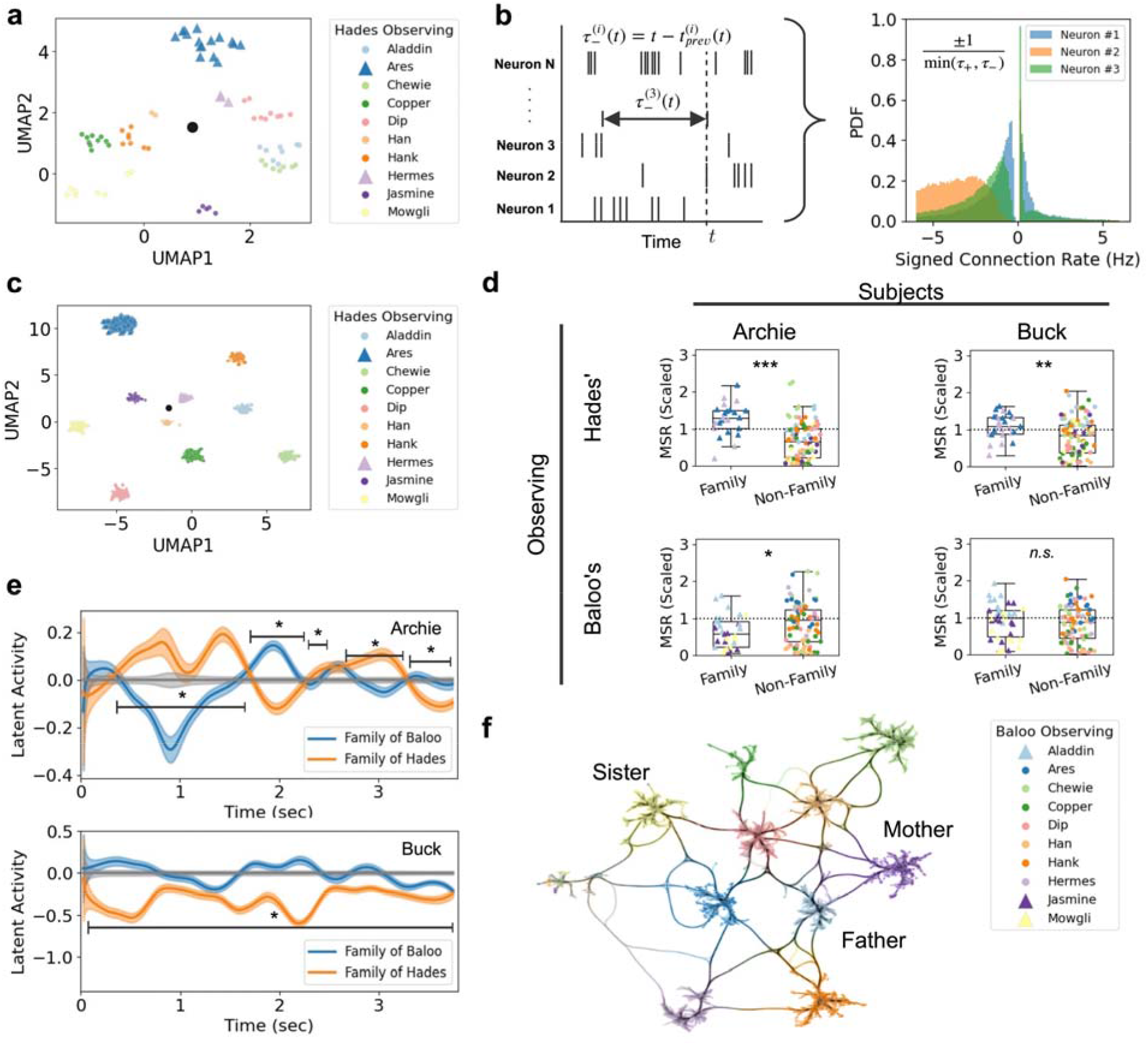
Cross-modal representation of identity using rate and event codes. **[a]** Two-dimensional manifold projection of our rate-coded representation computed from firing rates of identity-specific time bins. One symbol represents one identity match trial. Indicated is the mean (black). Colors in legend correspond to individuals. **[b]** Schematic illustrating the hindsight delay to a given neuron (left), used to generate histograms of signed connection rates to three neurons (right). **[c]** Two-dimensional manifold projection of our event-coded representation of identity computed as the manifold projection of signed connection rates of all neurons in the same exemplar recording session. One symbol represents one spike. Indicated is the mean (black). **[d]** Boxplots of MSR showing significantly different values when subjects observed family of other subjects. Shown is Archie observing family of Hades (top left, p<0.001, N_identities_≥23) and Buck observing family of Hades (top right, p=0.003, N_identities_≥26), Archie observing family of Baloo (bottom left, p=0.017, N_identities_≥30), and Buck observing family of Baloo (bottom right, p=0.828, N_identities_≥37). Significance was computed according to Student’s t-test. **[e]** Latent activity averaged over all recording sessions from subjects Archie (left) and Buck (right). Colors indicate average over the family of Baloo (blue) and Hades (orange) relative to all conspecifics (gray). Shaded regions indicate 95% confidence of the mean estimated via bootstrap. **[f]** Graph of connections bundled between individuals. Triangles in legend indicate family members as in [a,c,].

To investigate whether representation of identity can be described by the relative timing of spikes, we computed manifold projections of spike times recorded during match trials (Figure 4B, left) using parameterless *signed connection rate* features. The signed connection rate from one neuron to another describes how it interacts with other neurons, revealing statistical distributions specific to any given pair of neurons (Figure 4B, right); a facet of neural activity separate from the firing rate of any single neuron. Results using this event-coded measure again revealed excellent separability for identity-match trials (Figure 4C), thereby replicating the effect observed with the INM using an independent facet of neural activity and further supporting cross-modal representations of identity as encoded in population-level activity in marmoset hippocampus.

Given this result, we next asked whether social categories other than identity may likewise be represented in event-coded hippocampal activity. Specifically, we tested whether representations of other marmosets’ family members were distinct from non-family members for the two marmosets whose families were not included in the stimulus sets using two distinct quantifications of manifold projections, though the pattern was consistent for all subjects. First, results revealed a significant difference in the mean square range (MSR) of the manifold projections along this category boundary (Figure 4D, S12A), suggesting a larger event-coded state-space was occupied while observing family members (Figure S12B). Notably, while these projections were supervised, the clustering that emerged based on respective social relatedness was unsupervised. Second, we computed the unsupervised latent firing rate as the manifold projection of the absolute value of signed connection rate. Although individual identities did not separate (Figure S13A), we found trajectories that appeared stable in time and comparable across trials (Figure S13B). The motion of mean latent firing rate significantly separated social categories at multiple time points for all subjects (Figure 4E; Figure S13C,D). Together, these results demonstrate that neural representations of social identity in primate hippocampus are not only invariant to the sensory modality and comparable over time (Figure S14) but low-dimensional manifolds (Figure 4F) can describe relationships between different social categories (e.g. individual identity, family groups, etc.) learned by observing interactions between individuals (*8*).

Here we showed that the cross-modal identity of multiple conspecifics is represented in the primate hippocampus. Although we identified putative ‘concept cells’ similarly to humans (*5, 14*), we discovered that this population of highly selective neurons is not the only mechanism for representing concepts of individuals. Rather, both single neurons and the broader population in hippocampus encode cross-modal identity of multiple conspecifics, similar to what has been reported for objects (*23*), suggesting that the sparse representations of ‘concept cells’ may not be the only mechanism to represent semantic memory in hippocampus. Furthermore, analyses revealed that a population-level code represents not only the cross-modal identity of multiple familiar individuals but information pertinent to social categories, as well. Similar to the role of hippocampus in other contexts (*26*) (Figure S15), these representations may support a learned schema that here applies to social identity (*3, 4*). The presence of unimodal representations of identity in the primate frontal and temporal cortex (*1, 9*), amygdala (*11, 27*) and the medial temporal lobe (*28*) and representations of social dominance in amygdala (*29*) may reflect an integrative social recognition circuit in which substrates in the broader network play distinct but complementary roles that collectively govern natural primate social brain functions (*30*).

## Acknowledgements

We thank David Leopold for comments on a previous version of this manuscript.

## Funding

National Institutes of Health to R01 NS109294 (CTM)

National Institutes of Health to R01 DC012087 (CTM)

## Author Contributions

Conceptualization: CTM

Methodology: TT, MM, CTM

Investigation: MM, TT

Analysis: TT

Visualization: TT

Funding Acquisition: CTM

Supervision: CTM

Writing – Original Draft: TT, CTM

Writing – Review & Editing: TT, CTM

## The authors declare no competing interests

