## Supplementary material for "Cross-Modal Representation of Identity in Primate Hippocampus": Methods and Supplementary Figures

Science
AAAS

### Supplementary Materials for

#### Cross-Modal Representation of Identity in Primate Hippocampus

Timothy J Tyree[1,2], Michael Metke[1,3] & Cory T Miller [1,3*]

**This PDF file includes:**

Materials and Methods
Figs. S1 to S17

#### Materials and Methods

Subjects.

Four adult marmosets (2 male, 2 female) served as subjects in these experiments. All animals are socially housed with 2-8 conspecifics in the Cortical Systems and Behavior Laboratory at the University of California San Diego (UCSD). All animals housed in a cage are family members, as each cage comprises a pair-bonded adult male and female and 1-3 generations of offspring. The UCSD marmoset colony in the Miller Lab houses ~70 animals in 15 family groups in a single room with visual and acoustic access between cages. All procedures were approved by the Institutional Animal Care and Use Committee at the University of California San Diego and follow National Institutes of Health guidelines. A total of 47 recording sessions were performed with these subjects over the course of the experiment and analyzed here.

The total number of single units recorded from marmoset hippocampus totaled N=714 in Archie, N=822 in Baloo, N=212 in Buck, and N=610 in Hades (Figure S3B). All four subjects were considered equally in the identity neuron analysis and the MvMM neuron analysis (Figures 1, 2). All subjects were considered in the predictive time bin analysis (Figure 3) except for Buck due to his low count of single units across his 13 recording sessions. For the manifold projection analysis (Figure 4), all subjects were considered while they observed families that had at least two family members from amongst the cohort of individuals shown.

Experiment Design.

Neurophysiological recordings were performed while subjects were head and body restrained in our standard marmoset chair (*31*). Visual stimuli were presented on an LED screen from a BenQ monitor 1080 positioned 24 cm in front of the animal. Acoustic stimuli were presented at 70-80 dB SPL from a speaker positioned below the monitor (Figure S16). All behavior was collected in an anechoic chamber illuminated only by the screen, which had a dynamic range from 0.5 to 230 cd/$m^2$, with luminance linearity verified by photometer. Stimulus presentation was controlled using custom software and eye position was monitored by infrared camera tracking of the pupil. For hardware, calibration, and validation see previous work in the lab (*31*).

Subjects initiated trials by holding fixation of gaze for 100ms at a center fixation dot on the screen, at which point stimulus presentation was initiated. The 150ms period immediately post-stimulus was discarded to account for the time for visual signals to propagate from the retina to the hippocampus. This latency has been measured to be in the range 100-200ms (*32*). This biophysical argument supports our estimate of the stimulus onset t=0 occurring 150ms after stimulus was presented. Unless otherwise specified, baseline firing rates were estimated from 500ms preceding t=0 excluding 300ms for anticipatory firing. Stimulus responses were initially measured by comparing the peristimulus baseline firing rate to firing rates averaged from the max of a 500ms sliding window from t=300ms to 3.5s.

Stimuli were divided amongst unimodal– face-only and voice-only– and cross-modal– identity match and identity mismatch– on a trial-by-trial basis. Up to twelve conspecifics were represented per stimulus set (min 10, max 12). Face stimuli comprised multiple examples of each individual marmoset from different head orientation.

All face and voice stimuli were pictures or audio recordings from animals housed in the same colony room as the subjects. Because the colony is housed in a single room in which all animals have visual and acoustic interactions with each other, we assumed that all animals have sufficient experience observing each other to be familiar with their respective individual identities. Each individual marmoset was represented in multiple distinct stimuli ($N_{stimuli}$=36.0±15.3) for each individual in each recording session across each of the three stimulus classes: face forward, face profile and vocalization. No single stimulus was presented to subjects more than two times in a single test session. Monkeys with fewer than 10 presentations per individual in a recording session were not considered in any analysis. The stimulus duration of trials involving vocalizations (i.e. voice-only and cross-modal) necessarily varied because each "phee" call differed in duration (mean: 3.02±0.74s). The median face stimulus duration was 3.50 seconds (IQR: 2.78-3.51 seconds). The minimum face stimulus duration was 2.05 seconds and the maximum face stimulus duration was 4.46 seconds. Stimuli were presented in 10-trial blocks, with an inter-block active forage trial with juice reward to maintain attention. Each recording set was composed of 400 face and/or voice stimuli, split into 2 subsets.

All stimuli were composed of faces and/or voices of conspecific monkeys in our colony familiar to each subject. A total of 16 individual monkeys were represented overall (9 male, 7 female). Test subjects were not included in their own stimulus sets. Because our goal was to test for representations of individual identity rather than cross-modal perceptual integration of face/voice biomechanical movements (i.e. McGurk Effect) we presented subjects with static face stimuli so as not to introduce confounds that may emerge due to temporal misalignments of the face and vocalizations during the identity mismatch trials.

All face stimuli were photographs of monkeys from our colony taken while animals were in our standard marmoset chair with a light background behind them. The animals are trained to sit comfortably while a neck guard restricted their mobility. While seated, subjects could freely change head direction. Photographs of each subject were visually inspected and selected based on image quality and suitable representation of multiple head orientations (Figure 1A-C, S1). Photos used as stimuli were cropped to only show the neck guard and the face/head, so as to eliminate views of the rest of the body and chair.

All voice stimuli were marmoset "phee" calls comprising two pulses, the species-typical long-distance contact calls. Previous work has shown that marmosets are able to recognize the caller's identity when hearing "phee" calls (*33*). Recordings were made at 44.1kHz sampling rate while a monkey engaged in natural vocal interactions with a visually occluded conspecific in a soundproof chamber and hand-selected using custom code. Only examples with high SNR and minimal background noise were selected for stimuli.

All analyses were performed in Python unless otherwise indicated.

**Surgical and neural recording details.**

The surgical procedure employed here has been described previously (*16*). Briefly, we performed an initial surgery to affix a post to the skull on each animal to restrain subjects' head during experiment preparation. Following recovery, a second procedure was performed to embed the drive housing and the electrode array for stable chronic electrophysiological recording. We implanted a 64-channel microwire brush array (MBA, Microprobes) either unilaterally or bilaterally into the hippocampus using preoperative MRI stereotaxic coordinates. Electrode locations were confirmed by postoperative MRI and histology. All surgeries were performed under sterile and anesthetized conditions. The implants were inserted 7-13 degrees of angle off the vertical using the medial sulcus as reference before the operation has taken place. Neural recordings were performed with an Intan 512ch Recording Controller system via an RHD2164 64-channel amplifier chip, sampled at 30kHz. Neurophysiology data was analyzed using Spyking Circus yielding across all recording sessions 2,358 isolated units, referred to as neurons in the main text and in the remainder of Methods and Materials. Standard procedures were employed to remove obvious recording errors, which resulted in less than 1% of trials being removed from the analysis *a priori*.

**Identifying Identity Neurons.**

Hippocampal neurons were tested for an invariant response to individuals in the face-only and voice-only trials using an ROC analysis identical to that described in human hippocampus (*5*). For each isolated single neuron we performed the analysis for all identities where at least 4 unimodal stimuli (either face or voice but not both) were presented for each of the following three unimodal stimulus categories: face forward, face profile and voice.

The response of a neuron to a trial was the taken to be the maximum spike count in a 500 millisecond continuous sliding time window from t=0.3 seconds to t=3.5 seconds following stimulus onset at time t=0. As in *(5)*, the response of a neuron to a stimulus was the median response averaged over all presentations of the stimulus.

A neuron was considered responsive to a stimulus if its response to the stimulus was above the responsiveness threshold, which was determined as the sum of the mean baseline plus two standard deviations (s.d.) of the baseline, where the baseline was the number of spikes averaged over the times t=-0.8 seconds to t=-0.3 seconds. This differs from the original study in humans *(5)*, which used five s.d. instead of two, which was not practical in this study due to marmoset hippocampal neurons typically exhibiting larger baseline firing rates (Figure S2), for which five s.d. would have resulted in responsiveness thresholds that would only be evident in N=166 out of the 2,358 single units involved in this study (7.0%).

A neuron was considered cross-modal invariant to an individual if it was responsive to all three unimodal stimulus categories for that individual. If a neuron instead responded only to the voice of an individual, then it was considered voice-invariant. If a neuron instead responded to an individual for both the front facing and profile facing stimulus categories, then it was considered face-invariant.

As in *(5)*, stimuli were considered in ROC selectivity analyses only if at least one neuron responded to it. Also as in *(5),* an above-threshold response to a stimulus of the preferred subject was considered a positive test. Significance of an ROC for a given subject was determined by comparison to 99 surrogate ROC curves, which resulted from randomly and independently shuffling the labels. An area under the curve (AUC) that surpassed that of all surrogates was considered significant ($p<0.01$). Neurons that met or exceeded these thresholds were necessary to determine selectivity for individual identity in marmosets.

If a neuron was determined to be invariant to an individual within a given mode or modes, then selectivity was determined using the same mode or modes for that same individual. That is, cross-modal invariant neurons were tested for selectivity using all three unimodal stimulus categories, face-only invariant neurons were tested for selectivity using only front facing and profile facing unimodal stimuli, and voice-only invariant neurons were tested for selectivity using only the voice.

Cross-modal invariant neurons that passed the ROC selectivity test of *(5)* were considered selective for the identity and were thus labeled as putative “concept cells”. Because all voice-only unimodal stimuli were combined into a single stimulus category, voice-invariance would imply voice-selectivity for one identity if not for an additional statistical test that compared the median trial response to the voice stimuli of the preferred individual to that of all other individuals according to a one-sided Wilcoxon-Mann-Whitney test ($p<0.01$) with an above-threshold response constituting a positive prediction of the preferred individual. The comparable test was used to determine selectivity for the face-invariant neurons. The invariant neurons demonstrating selectivity were considered identity neurons.

**Identifying MvMM Neurons.**

Determination of MvMM neurons was achieved by comparing the median response of a neuron to identity match trials to the median response of that same neuron to identity mismatch trials. If a neuron was responsive to either match or mismatch trials, then a statistically significant difference computed according to a Wilcoxon-Mann-Whitney test qualified a neuron as a MvMM neuron ($p<0.05$). Preference of a MvMM neuron to match or mismatch trials was subsequently determined by a one-tailed Wilcoxon-Mann-Whitney test ($p<0.05$). Importantly, we did not preselect for neurons that were broadly stimulus driven, but focused analysis only during the median stimulus and compared activity between match and mismatch trials. This is reflected in the exemplar neurons selected for Figure 2. The match preferent neuron (Figure 2A) shows a difference in firing rate during presentation of the stimuli but is not broadly stimulus driven. By contrast, the mismatch preferent neuron (Figure 2B) exhibits stimulus driven activity as well as differential firing rate between the stimulus types.

**Identifying predictive time bins.**

Hippocampal neurons were analyzed in terms of their firing rate response during time bins that we identified as candidate time bins. For each neuron, our procedure consisted of three stages. The first stage was to generate a large list of time bins of varying duration using an extension of a sliding window approach. The second stage identified a subset of time bins as having a general ability to distinguish trials. We required this subset to be mutually disjoint. Candidate time bins resulted from the third stage, which varied each time bin independently according to our refining procedure.

The first stage extended the sliding window approach by using 200ms time bins evenly distributed between 0 and 3.6 sec, the maximum stimulus duration (Figure S4A). Time bins of duration greater than 200ms were constructed by joining adjacent time bins, leading to a maximum allowed time bin duration of 3.6 seconds. A general ability to weakly distinguish trials was determined by splitting the training trials according to three-fold stratified cross-validation and then computing the training AUC of each fold (Figure S4B). Training AUC was initially computed from the ROC curve that resulted from an above-threshold firing rate response determining a positive trial. Separately, training AUC was computed from a below-threshold firing rate response as determining a positive trial. In either case, if the training AUC was greater than chance (AUC>0.5) for all three folds, then the time bin was retained for stage two. The same convention for *above* versus *below* firing rate response as determining a positive trial was used for stage two and for stage three. All population-level decoders were blind to this convention of sign.

The second stage selected a disjoint set of candidate time bins, optimizing for their ability to distinguish trials by maximizing the mean AUC averaged over the same three folds. To achieve this, time bins were selected in decreasing order of their mean AUC and included only if doing so maintained the disjointness of time bins.

To reduce the effect of discretizing the trial into time bins, the third stage refined the resulting disjoint set by considering a number of random perturbations of each remaining candidate time bin and keeping only the optimal perturbation. The random perturbations shifted the start times and the end times independently by a random amount identically sampled from the normal distribution with zero mean and standard deviation equal to the duration of the unperturbed time bin. We generated a sample of N=100 perturbed time bins and removed those with a duration <10ms. Perturbations were additionally removed if they exhibited a start time before stimulus onset t=0 or if they exhibited an end time after t=3.6 seconds. A worsening AUC in any of the folds resulted in rejection of the given candidate time bin.

If any of the resulting training AUC values were smaller than that of the unperturbed time bin, there that perturbation was removed from consideration. The overall training AUC was computed for each perturbation using all training trials together. The perturbed time bin with the largest overall training AUC was kept instead of the unperturbed time bin. Perturbed time bins were allowed to overlap with other remaining time bins, thereby relaxing the condition of disjointness for the sake of parallelizability, which is statistically valid because zero spike times in the training set appear in the testing set and the decoder makes no assumption of independence of features. A flowchart summarizes the time bin refinement procedure (Figure S4C).

If no perturbations remained under consideration, then the unperturbed time bin was kept from stage two. Any remaining candidate time bins were considered predictive only if they presented a statistically significant difference in median firing rate for the true (e.g. identity match) training trials compared to the false (e.g. identity mismatch) training trials. Significance was determined according to $p<0.05$, where p was the statistic computed as the mean p-value resulting from a Wilcoxon–Mann–Whitney test conducted over the training trials averaged over five stratified cross-validation folds over training, which was a sufficient statistic in the sense that all time bins with $p<0.05$ also exhibited a statistically significant difference in median value at the same level of significance according to a Wilcoxon-Mann-Whitney test conducted over all MvMM trials. This procedure provided the features used in our population-level decoders. Data and code are made available to the reader (see Author Contributions).

**Training the population-level neural decoders.**

Population-level decoders were trained on the training trials before computing predictions for the separate testing trials. Decoders were trained and tested on a Quadro RTX 5000 GPU typically in less than five seconds of runtime.

The population-level decoders trained using firing rates directly as inputs. Neither translating nor scaling of the firing rates was performed, as the decoders were both location and scale invariant (*22*). The prediction was estimated by the weighted average of values returned by an ensemble of gradient-boosted decision trees relative to a default value of one half (controlled by base_score in Table 1). For each training epoch, at least 25 decision trees were trained (controlled by num_parallel_tree). While a unique solution exists for a given decision tree, a heuristic algorithm was used to approximate the unique solution using the quantile method of (*34*).

Decision trees were trained to minimize the binary cross-entropy loss function (equivalently, to maximize likelihood) at the ensemble-level by considering only a fraction of the training trials (controlled by subsample). Decision node rules considered only a fraction of the input firing rates (controlled by colsample_bynode) to determine placement of its weight. The weight of a node was limited to a certain amount (controlled by max_delta_step). The complexity of the decision node rules was further limited using linear and quadratic regularization (controlled by reg_alpha and reg_lambda in Table 1, respectively).

Each decision tree was gradient boosted in the sense that nodes were recursively added in accordance with an estimate of the gradient of a training loss computed at the ensemble-level. If inserting a decision node failed to improve the loss by a sufficiently large amount (controlled by gamma), then that decision node was removed from the tree. To further limit structural complexity, the maximum tree depth was set to no more than five decisions (controlled by max_depth). The weight for a new decision tree was scaled down by a factor (controlled by learning_rate). Training terminated for a given decision tree when the total weight for the next decision node was smaller than a certain amount (controlled by min_child_weight). After all decision trees terminated training, the training epoch was complete. After a fixed, predetermined number of training epochs, the ensemble terminated training. Then, predictions were computed for the testing trials (Figure S7A). Predictions were used to evaluate the predictive ability of a given set of one or more predictive time bins in terms of AUC.

**Determining hyperparameter settings for the population-level neural decoders.**

The parameter settings for our population-level neural decoders resulted from a series of coarse grid searches each conducted over a wide range of settings for one pair of hyperparameters at a time. Each parameter setting considered five-fold stratified cross-validation involving the training trials only with the goal of maximizing mean testing AUC. Early stopping was used during this tuning procedure, which supported a minimum 60 training epochs for the match vs mismatch (MvMM) predictive population and a minimum 67 training epochs for the identity-specific predictive population as sufficient according to early stopping. By increasing the number of training epochs, stability of performance became immediately apparent for up to 500 epochs for both MvMM and identity-specific decoders. We made no use of early stopping anywhere else apart from the hyperparameter tuning procedure described here.

This hyperparameter tuning procedure was conducted only on the training trials for Archie observing Waylon in one recording session from subject, Archie (session #8). Archie (male) and Waylon (female) were not family members– though they likely knew each other in the colony. These training trials (from session #8) were complementary to testing trials from no more than one of the multiple recording sessions summarized in Figure 3. The hyperparameter settings that resulted are reported in Table 1.

| **Hyperparameter** | **MvMM** | **identity-specific** |
|---|---|---|
| base_score | 0.5 | 0.5 |
| num_parallel_tree | 25 | 50 |
| subsample | 0.2 | 0.2 |
| colsample_bynode | 0.1 | 0.1 |
| max_delta_step | 0.5 | 1 |
| reg_alpha | 0.4 | 0.3 |
| reg_lambda | 0.4 | 0.3 |
| gamma | 0.1 | 5 |
| max_depth | 5 | 2 |
| learning_rate | 0.9 | 0.6 |
| min_child_weight | 0.5 | 1 |

**Table 1** Table of hyperparameters for our population-level neural decoders. Numerical values were passed as keyword arguments to the constructor of xgboost.XGBClassifier instances *(22)*. Columns correspond to the two types of predictive populations reported in the main text.

**Summarizing testing performance from multiple predictors.**

Population-level decoders were trained as MvMM or identity-specific predictors for each individual identity in each recording session involved in Figure 3. To account for variations in prediction magnitude between decoders, predictions were scaled linearly to a maximum value of unity before combining ROC traces in the multiple recording sessions summarized in Figures 3G-I,K and S9-10. No such scaling was involved with the multiclass predictions reported in Figures 3J and S8.

**Sampling trials for multiple predictive populations from the same recording session.**

For a given recording session, the following criteria were respected while partitioning testing trials from training trials involving the identity network model (INM) discussed in the main text. Testing trials for the INM were also testing trials for both the MvMM decoder and the identity-specific decoders. Because stimuli involving individuals were sampled uniformly, the frequency of a given individual could be small for a given recording session. To account for this, individuals were considered only if they exhibited at least forty appearances in a given recording session.

Because of the uniform nature of our uniform random sampling of trials over the larger space of cross-modal stimuli, each recording session had relatively few trials involving both the face and the voice of a particular individual. This resulted in far more negative trials being presented to the observer relative to the number of true trials for the INM. This was also the case for both the MvMM decoders and the identity-specific decoders reported in Figures 3 and S8-10. All three binary classification tasks had balanced samples randomly selected, which were then randomly shuffled before 30% were randomly selected to be testing trials. The remaining 70% of trials were considered for training. Unbalanced sampling in the training set was accounted for by scaling the positive weights by a factor of 5 for the MvMM decoders and 100 for the identity-specific decoders. Decoders involved in Figure 3 used 200 training epochs, all of which were used in testing decoder performance except the first training epoch. The only exception was the identity-specific decoders involved in evaluating the INM for the winner-take-all model in Figures 3J and S8, which considered all 500 training epochs.

**Decoding multiple identities using a winner-take-all model.**
We used the winner-take-all model to predict the identities of multiple individuals shown during identity match and face-only trials. The twelve individuals summarized (Figure 3J) have their detailed testing performance reported (Figure S8). The winner-take-all model predicted the correct identity with an overall testing accuracy of 91.0% ($N_{trials}$=454). For a given recording session, the following procedure was performed to generate the predictions for the winner-take-all model. First, we identified all identities involved in a sufficient number of identity match trials ($N_{trials} \geq 12$). All identity match trials involving the identities identified were shuffled and 30% were randomly selected as testing trials to be withheld from training with the remaining 70% of trials.

We considered predictions of our INM to approximate a predicted probability that a given trial from the testing set involved the given identity. The presence of the individual was modeled using the decoder outputs in the winner-take-all model if the INM had the sufficient number of predictive time bins available. After repeating this procedure for all individuals in the recording session, the predicted identity of the winner-take-all model corresponded to that of the maximum predicted value (Figure S7B).

**Quantifying relative contribution of identity neurons in decoders of preferred identities.**
To investigate the possibility of identity neurons exhibiting any clearly observable significance in the INM at the population-level, we removed all identity neurons from consideration and recomputed the testing predictions of Figure 3I for each individual that was statistically preferred by an identity neuron. After recording the testing AUC, we repeated a comparable procedure as a control that randomly removed an equivalent number of predictive time bins from any neuron that was not found to be an identity neuron. This control procedure was repeated many times ($N_{samples}$=200) and then averaged to estimate the mean control testing AUC, which was not significantly different from a normal distribution according to D'Agostino-Pearson's omnibus test ($p>0.05$, $N_{samples}$=200). The aforementioned control and test procedures were conducted using independent randomized samples.

ROC curves were computed with above-threshold values indicating a positive trial for the three observers with at least two family members amongst the identities presented. The INM appeared successful despite the removal of identity neurons independently for multiple observers (Figure S10). Removing identity neurons from the INM for all recording sessions involving one observer resulted in a mean testing AUC that was not significantly smaller than that of the control according to a one-tailed paired student's t-test that supposed identity neurons contributed more to decoding than other neurons. We independently replicated this same statistical insignificance of identity neurons at the population-level for multiple observer subjects ($p>0.05$, $N_{observers}$=3). This insignificance was consistent with a comparable analysis that made no assumption of normality, which suggested the median testing AUC was also not significantly smaller when all identity neurons were removed relative to the control ($p>0.05$, $N_{observers}$=3). It is uncertain whether this insignificance can be attributed to these identity neurons being observed in nonhuman primates, as no comparable predictive time bin analysis has ever been performed in humans to the knowledge of the authors.

**Computing signed connection rate.**

Our event-coded representation relied on our signed connection rate measure, which we computed using our two primitive event measures. The first we referred to as the hindsight delay, $\tau_- > 0$, which is the amount of time since a given neuron has spiked. The second we refer to as the foresight delay, $\tau_+ > 0$, which is the amount of time until a given neuron will spike. A schematic illustrating the computation of the hindsight delay is shown (Figure 4B, left). A similar computation is found for the foresight delay by time inversion. If the given neuron has not yet spiked, then we take the hindsight delay to approach infinity. Similarly, if the given neuron was not observed to spike again, then we take the foresight delay to approach infinity. Note that our primitive event measures do not evaluate to non-positive real numbers.

The magnitude of our signed connection rate is the multiplicative inverse of the minimum of the hindsight delay and the foresight delay. Finally, we set the sign of our signed connection rate to be negative if the hindsight delay was used. Using the standard conventions of real analysis, our signed connection rate is now well-defined at all times for all neurons that exhibited at least two spikes. Equivalently, our signed connection rate was computed according to a real function of two variables

$$c(\tau_+, \tau_-) = \frac{\Theta(\tau_- - \tau_+)}{\tau_+} - \frac{\Theta(\tau_+ - \tau_-)}{\tau_-},$$

where $\Theta$(x)=1 if x is nonnegative, otherwise, $\Theta$(x)=0.

We evaluated our signed connection rate for every neuron at the spike times of the neuron that spikes the most over the recording session (i.e. the reference neuron). This was our attempt to measure how a single neuron “connects” with any other neuron. In doing this, we observed statistical distributions that appeared specific to a given neuron pair (Figure 4B, right). We considered a given neuron to have an approximately symmetric signed connection rate if it exhibited no more than twice as many negative values as positive values in these statistical distributions.

**Estimating manifold projections.**

We used uniform manifold approximation and projection (UMAP) to compute our manifold projections in Figure 4 of the main text, which presents descriptive manifold projections computed from predictive firing rate features and separately from our signed connection rate measure of spiking events. The same parameter settings on the same optimization algorithm was used for both rate and event-coded manifold projections. We used the identity-specific predictive time bins in the rate-coded representation. The rate-coded manifold projections considered neuron spikes from t=0 to 2 seconds after the stimulus onset. Similarly, the event-coded manifold projections considered neuron spikes from t=0 to 2 seconds after the stimulus onset. The average predictive time bin from the MvMM predictive population reported in Figure 2 was centered from t=0 to 2 seconds after the stimulus onset, with approximately half of predictive time bins ending earlier, which supports 2 seconds as a reasonable choice for the max time considered by the rate and event-coded manifold projections.

The UMAP algorithm was composed of two steps that can fruitfully be described as graph construction and graph projection (*24*). The graph was constructed from a given set of comparable observations. The graph was projected to a low-dimensional space of real numbers. The output was embedded in twenty-four-dimensional real space for statistical analyses and two to three dimensions for visualizations. In the optimization procedure, five negative samples were selected for each positive sample. The minimum distance between two observations was set to 0.1Hz. The number of nearest neighbors was initialized to 50 for rate-coded representations and 1000 for our event-coded representations. Repulsion strength was initialized to unity. Local connectivity was set to 1Hz in estimating probability distances. We trained for 200 epochs at a learning rate initialized to unity. The resulting function was equipped with a learned graph of the data, which projected to the manifolds visualized in Figures 4, S11, and S14-15. An example of connections from such a learned graph were visualized (Figure 4F).

For our rate-coded manifold projections, the inclusion of predictive time bins ($p<0.05$) appeared sufficient for the separation of individuals (Figure S11A), which was supported by computing the minimum distance between the centroid of any individual and then comparing across multiple recording sessions. Minimum distances that were computed from predictive time bins exhibited a significantly smaller median value when compared to candidate time bins that were not predictive ($p>0.85$) according to a Wilcoxon-Mann-Whitney test ($p<0.001$, $N_{sessions}=29$), suggesting predictive activity leads to better separation of individuals in comparable rate-coded representations (Figure S11B). Shown are examples of rate-coded manifold projections that used predictive firing rates as trial-by-trial observations. Event-coded manifold projections used signed connection rates as spike-by-spike observations for Hades (Figure S11C,D) and for Baloo (Figure S11E,F) in addition to Archie (Figure S14A-C) and Buck (Figure S14D-F).

**Estimating latent firing rate.**

Our latent firing rate was computed using unsupervised nonlinear dimensionality reduction of the absolute value of the signed connection rate for all neurons that had no less than one third of its computed signed connection rate values as positive (i.e. approximately symmetric). In computing the latent firing rate, we used a method of nonlinear dimensionality reduction that made no assumption of uniformity, which was achieved by passing the keyword argument, densmap=True to the manifold projection constructor, umap.UMAP, in the Python programming language. The output metric and the input metric were both Euclidean (flat), which supports the output having the same units as the input. The output was embedded in six-dimensional real space and the first three dimensions are visualized in Figure S13A for an exemplar recording session. After this output was computed at the spike times of all neurons involved, it was analyzed as a time series by time ordering the data according to evaluation time.

By considering latent firing rates evaluated at the times t=0 to 4 seconds after a stimulus onset, we observed relatively stable trajectories for multiple recording sessions conducted over multiple observers. Shown are three exemplar identity match trials, where Baloo observed the face and voice of her mother, her father, and her sister as shown in Figure S13B. We performed a median filter with a sliding window of 50 neuron spikes before plotting our estimates of the latent firing rates.

**Generating the hammer bundle plot.**

The graph of connections bundled between individuals in Figure 4F represents the learned graph associated with an event-coded representation of identity analogous to Figure 4C. The procedure for generating the shape of Figure 4F was achieved using the Python function, umap.plot.connectivity with the keyword argument, edge_bundling='hammer'. Coloration was achieved to multiplying the resulting image with a color mask. The color mask resulted from passing the colored scatter plot of the event-coded representation through a Gaussian filter using the GNU Image Manipulation Program, which was also used for the image multiplication.

**Determining anatomical positions of implants.**

All implants were followed by at least one postoperative MRI (Figure S17). The scans were aligned to anatomical features with RadiAnt Dicom viewer and the position along the anterior-posterior axis was determined by measurement from the center of the array to the ear canal. Because implants were stereotactically performed coronally, all recordings for a given array were assigned the same anterior-posterior (AP) position.

Because of the 1mm spread of the microwire brush arrays, it was difficult to precisely estimate the position of any given electrode, or indeed the entire bundle on a particular day. We used the position of the tip of the electrode from each MRI and extrapolated the trajectory by estimating position along the drive axis by cross-referencing with contemporaneous notes made of the date and distance of every movement of the drive. Based on a centroid at each estimated position, we chose particular sessions for we had the greatest confidence that the majority of the array was located predominantly in one or two hippocampal fields. Because the relative positioning of individual electrodes was not clearly observable, all reported analyses were developed to be agnostic to neuron location.

**Confirming implant location by MRI.**

MRI was performed at the UCSD Center for Functional Magnetic Resonance Imaging in a 7.0T Bruker 20cm small animal imaging system using Advance II software. Preoperative images were analyzed in Osirix DICOM Viewer and stereotactic coordinates were established using a pair of saline-filled barrels affixed above the putative posterior end of temporal sulcus (marked on the skull during headcap surgery). Array positioning and tract trajectory was verified by post-operative MRI. Follow-up scans were performed occasionally to update array position.

Determination of anatomical positioning was performed using RadiAnt DICOM Viewer (Medixant, n.d.). Stereotactic alignment was performed using a number of clearly defined and readily identifiable anatomical landmarks. 2D coronal slices were made vertical by rotating to align the medial longitudinal fissure with a vertical line. Yaw was corrected by re-slicing the coronal plane to align both interaural canals. Pitch correction was performed by re-slicing MRI so that the $4^{th}$ ventricle was aligned vertically with the isthmus of the corpus callosum.

Position on the anterior-posterior axis was calculated relative to the interaural canal. Measurement was taken from the coronal slice at which the array first entered the hippocampal complex (Figure 1D, 2E). Arrays were implanted with as little pitch as possible, so AP position variability is negligible along the electrode trajectory.

Electrode positions are not precisely determinable with our brush arrays, as microwires are not visible at the resolution of the scans and individual tips are not individually distinguishable by any practical means available. Electrode splay of the 64-ch MBA in tissue was measured at approximately 1mm, so we approximated electrode position by use of a 1mm spherical voxel centered at the tip of the array.

We used a Microdrive with a 500µm thread pitch that could reliably make controlled movements with a precision of 30-40µm. An array tip was identified for every MRI in each subject and position was extrapolated based on contemporaneous notes regarding electrode movement. Once putative array centroids have been hand-tagged they were assigned to one of the hippocampal subfields. Centroids were deemed to be in a hippocampal subfield if more than 70% of their volume fell within that area, as assessed by hand-traced MRI. Recording sessions where the centroid fell significantly between two subregions were not counted in anatomical analyses. CA2 and CA3 were combined due to insufficient granularity in this methodology and resolution in our scans to effectively differentiate them. Figure S6 shows the estimated position of each electrode array in the hippocampus for all subjects.

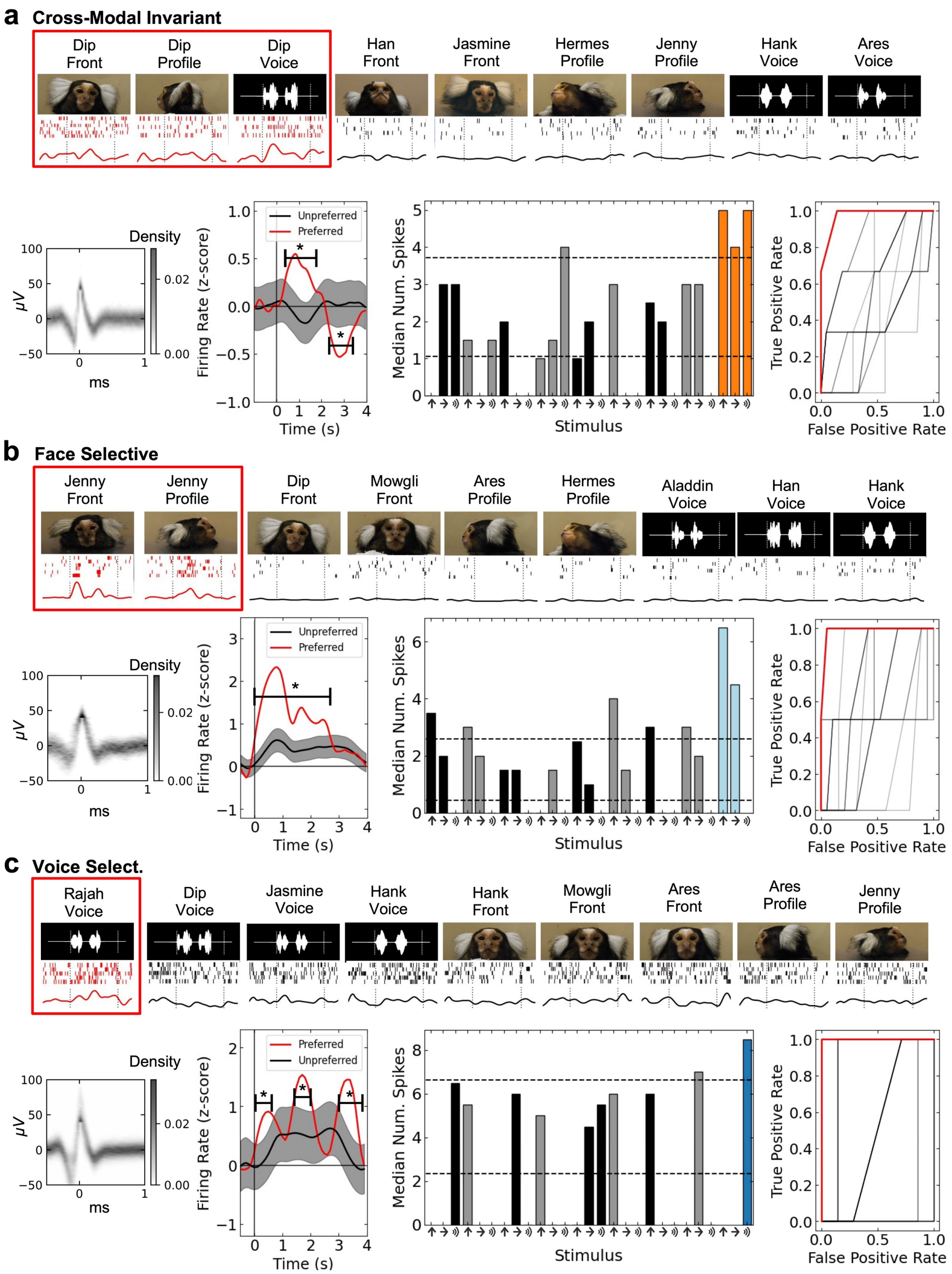


**Figure S1 Supplementary Exemplars** Shown are exemplar identity neurons that are **[a]** cross-modal invariant**, [b]** face-selective, and **[c]** voice-selective comparable to Figure 1A-C. **[a-c]** Top row: subset of stimuli shown above raster and peristimulus time histogram (PSTH). Bottom row: spike waveform; normalized PSTH to all stimuli (preferred: red, nonpreferred: black), indicated are time points that show significant difference (p<0.05); median number of spikes for unimodal stimuli (grey/black indicate non-preferred individuals; ROC curve (shuffled controls shown in black). PSTH was normalized by the pre-stimulus baseline, and shaded regions indicate 95% confidence intervals. Indicated are time points that show a statistically significant difference in mean (p<0.05). Horizontal dotted lines indicate mean background firing rate and responsiveness threshold.

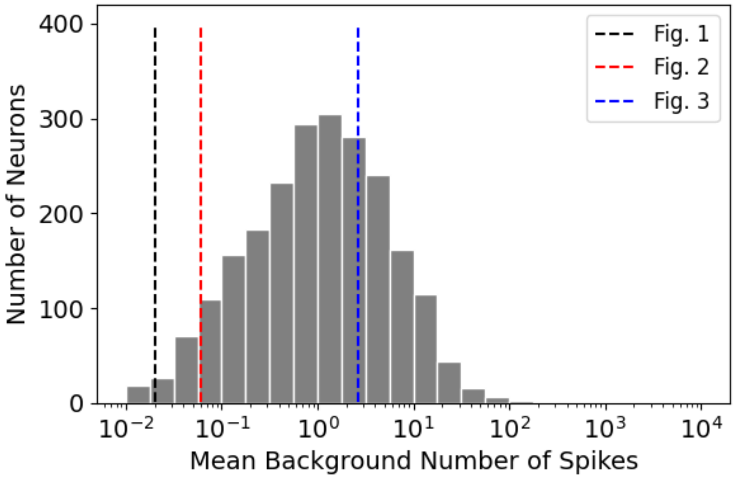


**Figure S2 Marmoset hippocampus neurons have high baseline firing rates.** Shown is a histogram of the mean background spike counts computed for all neurons involved in this study. The dotted lines come from the mean baselines reported in the main figures from Quian Quiroga *et al.*, *Nature* (2005), which summed over 700ms instead of 500ms.

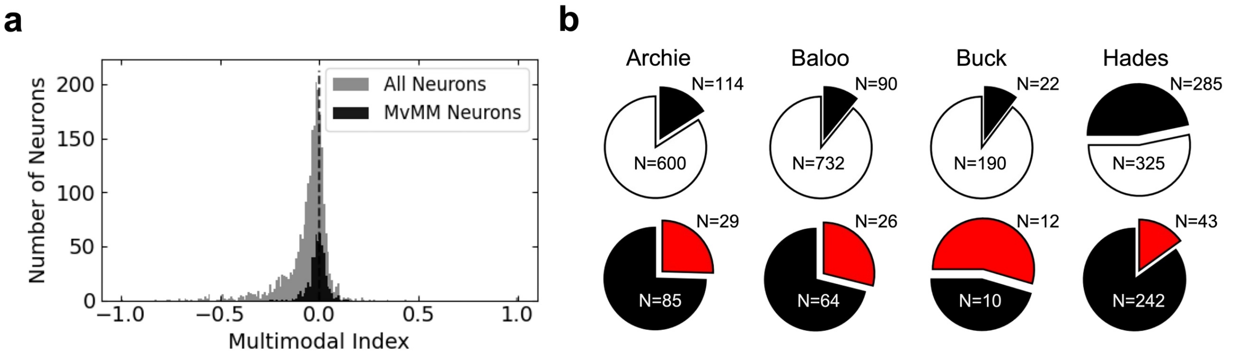


**Figure S3 [a]** Histogram showing the multimodal index of MvMM neurons (black) and all recorded neurons (gray). Neither the mean nor median multimodal index was significantly greater than zero for either population ($p>0.05$, $N\geq499$). The multimodal index was not well defined for N=12 out of 511 MvMM neurons due to small response. Zero is indicated by the black dotted line. Bin width is 0.01. **[b]** Pie charts showing the abundance of MvMM neurons averaged over all recording sessions for each observer involved in this study. Shown is the number of MvMM neurons (black, top) amongst all other recorded neurons (white, top) and the number of identity-match preferring MvMM neurons (black, bottom) amongst the identity-mismatch preferring MvMM neurons (red, bottom). The CA1 region was only confirmed in $N_{sessions}=4$ out of 8 of the recording sessions from Hades.

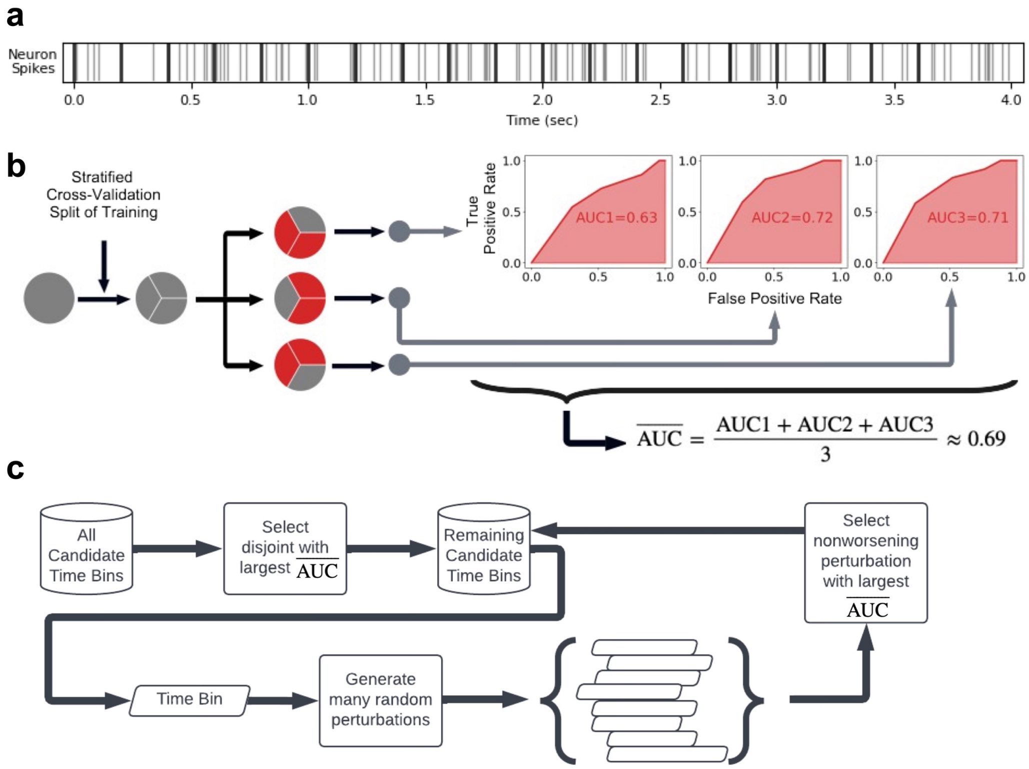


**Figure S4. Identification of predictive time bins. [a]** Schematic showing (gray) the spike times of an example neuron firing versus time after the stimulus onset at t=0. Indicated are (black) start and end times of time bins before the refinement procedure. **[b]** Flow chart showing training trials being split by stratified cross-validation to result in multiple receiver operator characteristic (ROC) traces. Each training fold resulted in an area under the curve (AUC), which were then averaged to produce the mean training AUC as an estimator of the general ability of a time bin to distinguish true trials from false trials. Time bins satisfying a list of properties were considered as candidate time bins (described in Methods). **[c]** Flow chart showing the procedure that resulted in all predictive time bins (described in Methods).

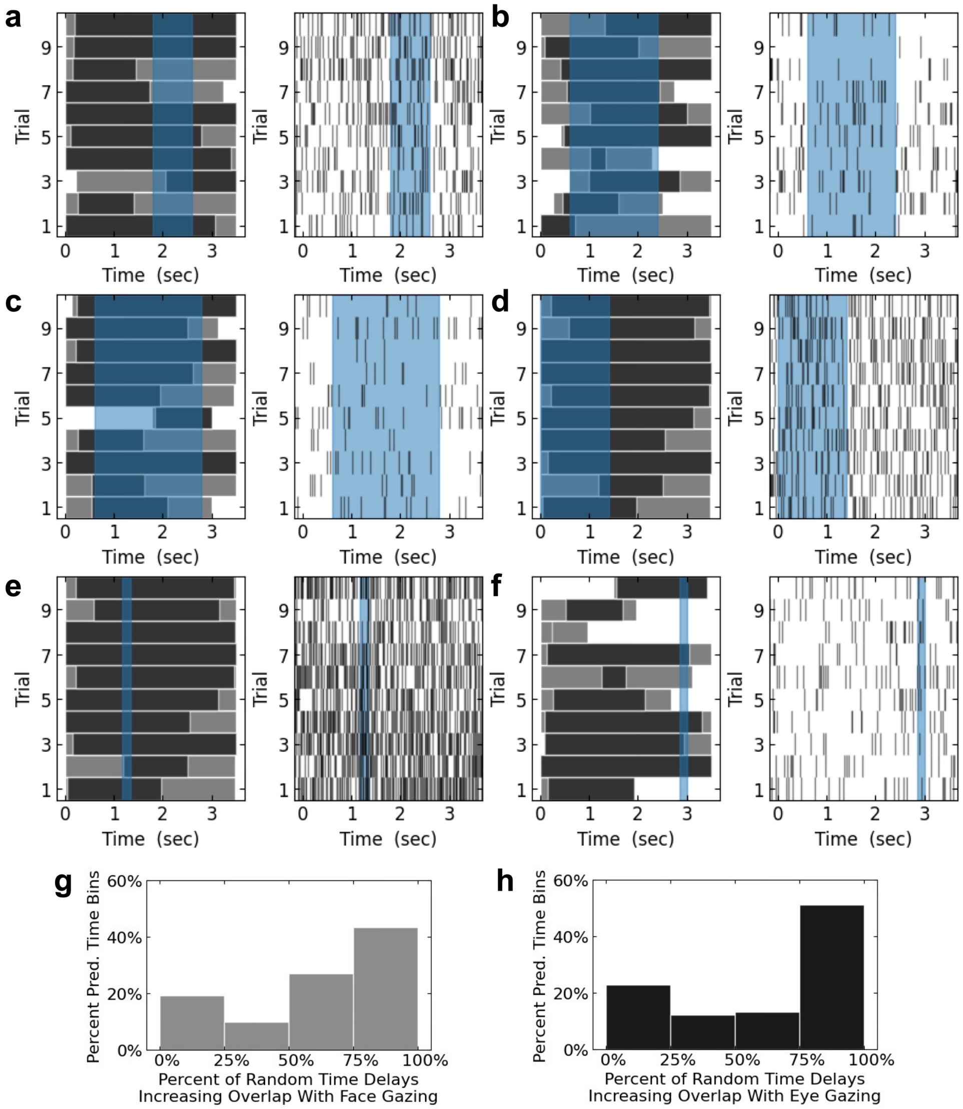


**Figure S5 Variability of visual behavior relative to identity-specific predictive time bins [a-f]** Shown are visual behavior rasters (left) and spike rasters (right) for six predictive time bins. Blue shaded regions indicate the identity-specific predictive time bin. Gray indicates face gazing while black indicates eye gazing in the visual behavior rasters. Trials represent repeated presentations of the same front-facing unimodal stimulus. Unimodal stimuli were chosen to agree with the identity preference of the predictive time bin. **[g-h]** Histograms showing the relative abundance of random delays that increased the amount time in common between the time bin and time spent gazing at preferred faces (g, gray) and time spent gazing at preferred eyes (h, black). Bar height shows the percent of identity-specific predictive time bins, where each time bin had at least 10 presentations of at the same unimodal face-only stimulus where both eyes of the preferred individual were clearly visible ($N_{bins}$=218). More area in the right two bars indicates perturbing the time bins typically decreased overlap with visual behavior.

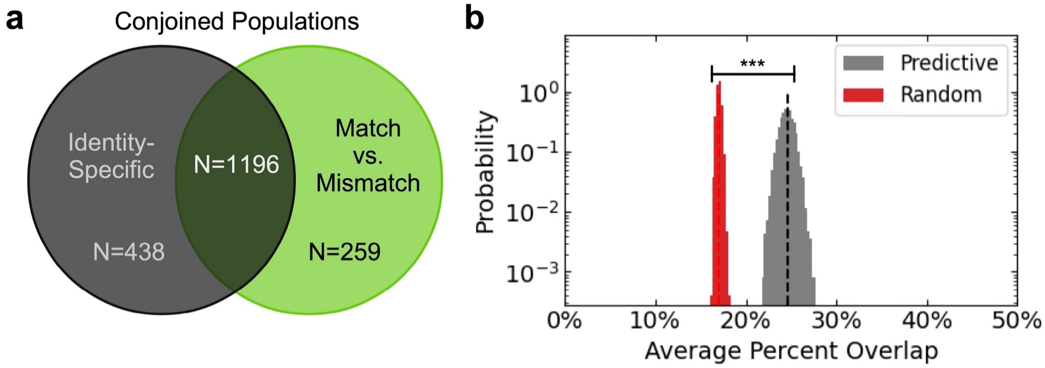


**Figure S6 [a]** Venn diagram showing the abundance of predictive neurons in common between the identity-specific predictive neurons (black) and the MvMM predictive neurons (green). **[b]** Histograms showing the probability density of the average percent overlap of the identity-specific predictive time bins with the MvMM predictive time bins from the same neurons (gray) and of an equal number of uniformly distributed pairs of random time bins as control (red). Indicated is the total duration of overlap divided by the total duration of identity-specific predictive time bins, (612.3s/2493.7s)=24.6%±1.5% (black dashed line), which was significantly greater than control (17.0%±0.5%; red dashed line) according to Student's t-test ($p<0.001$, $N_{samples}=10,000$). Control uniformly sampled pairs of time bins on the interval from t=0 to t=3.5 seconds following stimulus onset. The bin width is 0.25%.

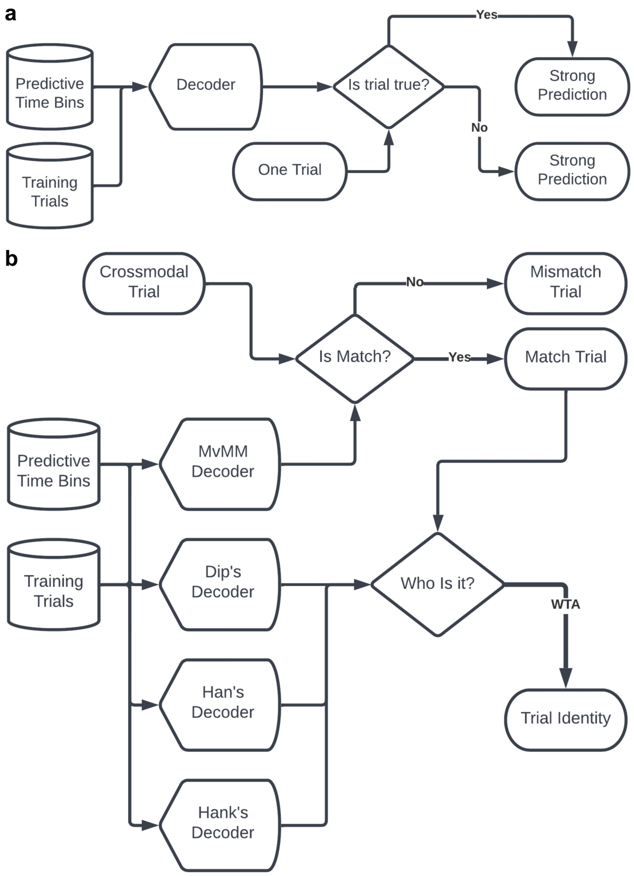


**Figure S7. Decoder Schematics [a]** Flow chart showing predictive time bins were combined with the training trials that were used to determine the predictive time bins to train a decoder for classifying trials as either true or false. The decoder then produced remarkably strong predictions on novel trials. **[b]** Flow chart showing the winner-take-all model resulting from a MvMM decoder and one identity-specific decoder for each individual. Cross-modal trials were categorized as either match or mismatch trials. The identity of the match trial was then predicted as that of the decoder with the largest output via winner-take-all (WTA).

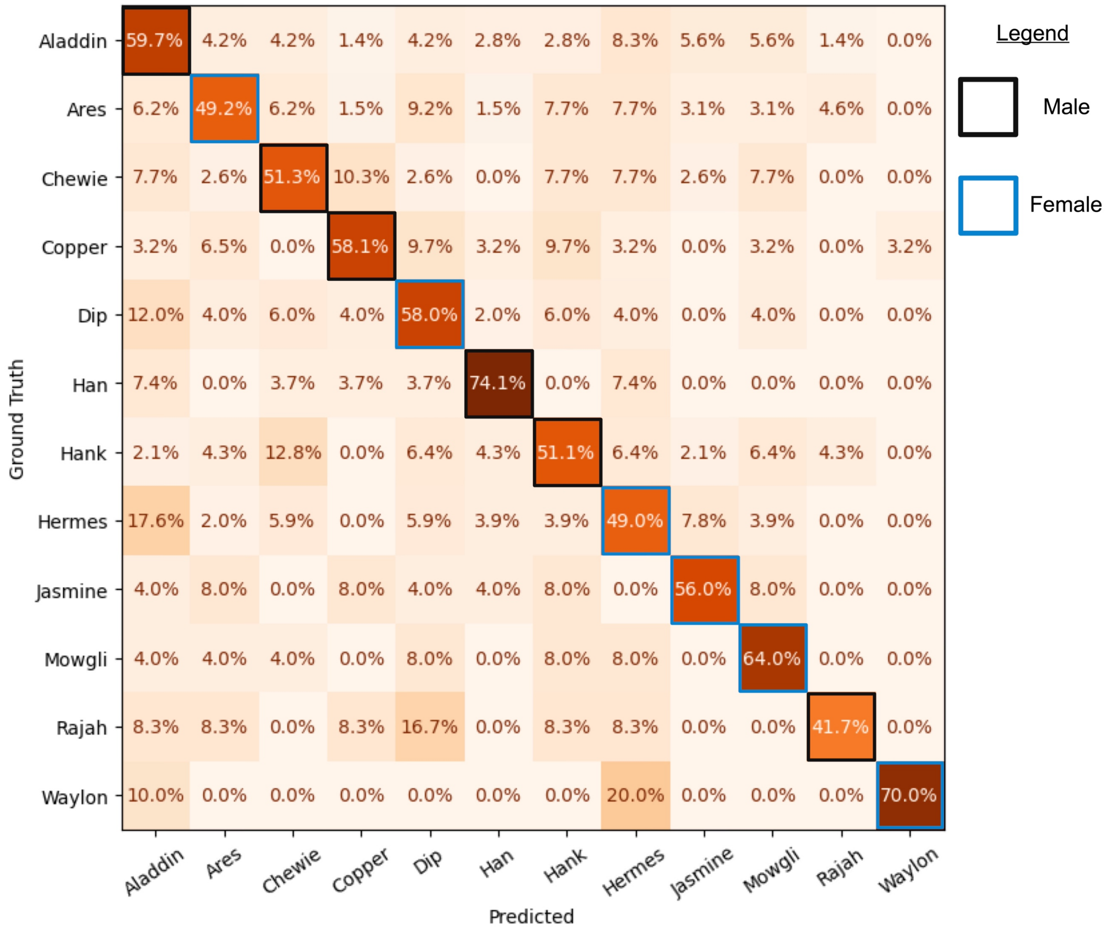


**Figure S8. Multiple individuals classified by winner-take-all model.** Confusion matrix reporting the winner-take-all predictions of the INM on twelve individuals shown to three observers over 34 recording sessions (testing accuracy=0.91, sensitivity=0.91, specificity=0.91, precision=0.88, negative predictive value=0.93, $N_{trials}$=454 match trials). The biological sex of the observed conspecifics is indicated by on the diagonal with blue indicating female and black indicating male. The following conspecifics were family members with a subject: Aladdin, Jasmine, Mowgli, Ares, Hermes. Percentages indicate true positive rates of the testing set of trials. All individuals decoded testing trials with a true positive rate at least 5X random chance, as is indicated by the black dashed line in Figure 3J of the main text.

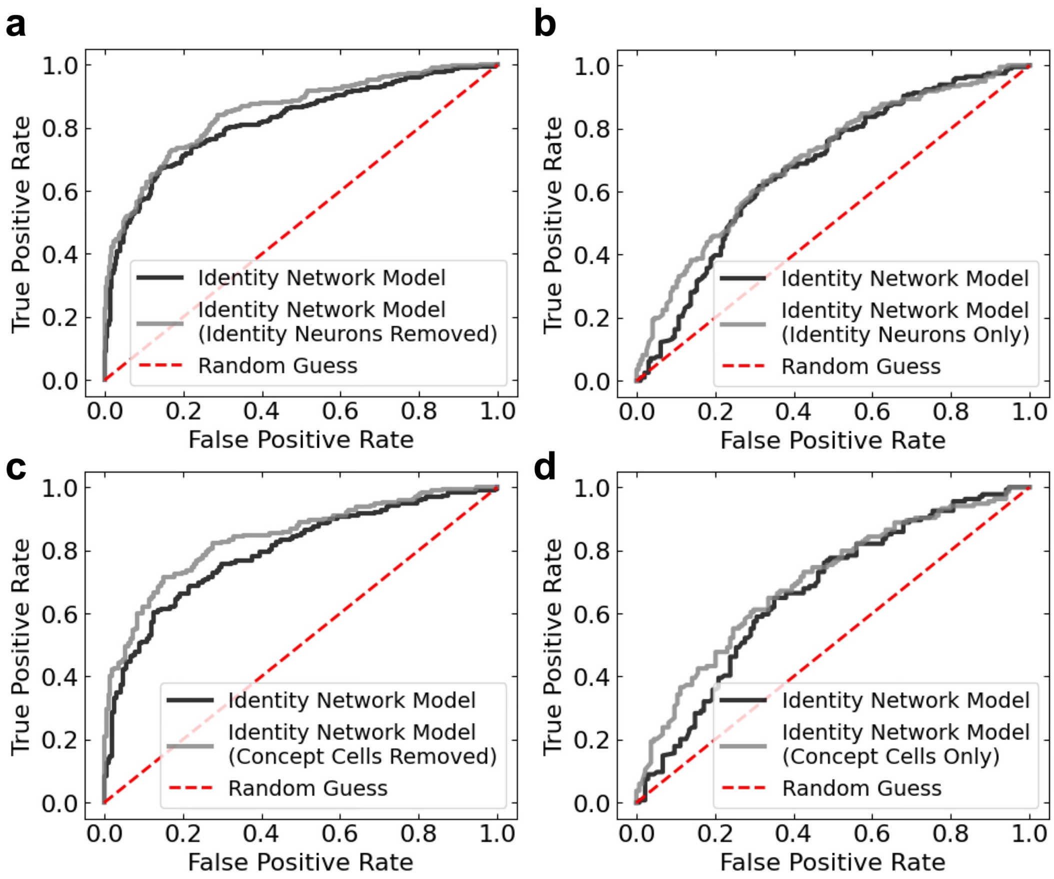


**Figure S9. Decoding performance with and without identity selective neurons averaged over preferred individuals. [a]** Shown are the ROC traces of the INM with all identity neurons removed (gray; AUC=0.850) and an equal number of random neurons removed from the remaining predictive population (black; AUC=0.820). **[b]** Shown are the ROC traces of the INM with only identity neurons considered (gray; AUC=0.700) and an equal number of neurons randomly selected from the remaining predictive population as control (black; AUC=0.677). Indicated is random chance (red dotted; AUC=0.500). **[c]** Shown are the ROC traces of the INM with all putative "concept cells" removed (gray; AUC=0.841) and an equal number of random neurons removed from the remaining predictive population (black; AUC=0.795). **[d]** Shown are the ROC traces of the INM with only putative "concept cells" considered (gray; AUC=0.700) and an equal number of neurons randomly selected from the remaining predictive population as control (black; AUC=0.666). Indicated is random chance (red dotted; AUC=0.500).

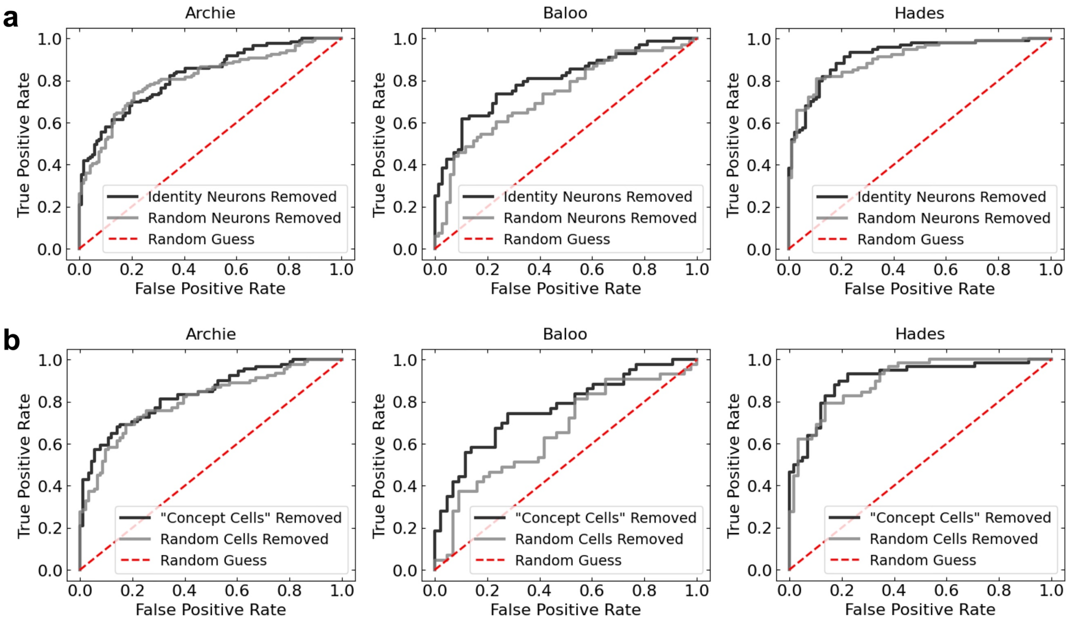


**Figure S10. Identity network model for individual subjects.** ROC curves were computed by averaging over all recording sessions for each of three observers: Archie (left, $N_{sessions}$=14), Baloo (middle, $N_{sessions}$=12), and Hades (right, $N_{sessions}$=8). **[a]** ROC curves of our INM with only identity neurons (black) and an equal number of cells from the remaining predictive population (gray). Individual identities were averaged over if they were preferred by at least one identity neuron. **[b]** ROC curves demonstrating the predictive power of our INM with all "concept cells" removed (black) and an equal number of cells removed from the remaining predictive population (gray). Individual identities were averaged over if they were preferred by at least one "concept cell". We controlled for network size by using the same number of features for both ROC curves in each panel. We did this for both the MvMM predictive population and the identity-specific predictive population in evaluating the INM.

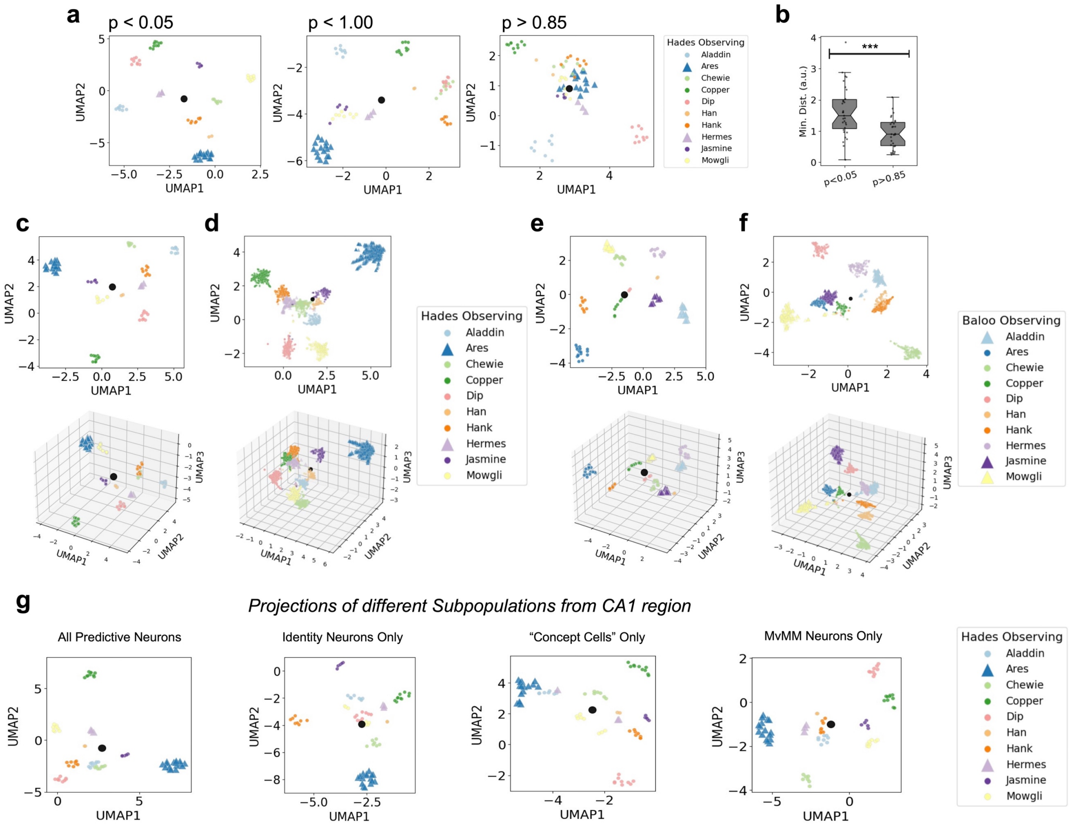


**Figure S11. Low-dimensional projections of our rate code and event code. [a]** Scatter plot showing an exemplar recording session as two-dimensional rate-coded representations of individual identity, where the firing rates were computed from all candidate time bins exhibiting (left) $p<0.05$, (middle) $p<1.00$, and (right) $p>0.85$. **[b]** Box-and-whisker plots showing the minimum distance between any individual in our rate-coded representation of individual identity. The median minimum distance of (left) $p<0.05$ was significantly smaller than the median minimum distance of (right) $p>0.85$ according to a Wilcoxon-Mann-Whitney test ($p<0.001$, $N_{sessions}=29$). **[c-f]** Shown are the (top) first two axes and (bottom) first three axes of our representations of individual identity for two distinct observers: **[c,d]** Hades and **[e,f]** Baloo. **[c,e]** Shown are manifold projections of our predictive time bins and **[d,f]** our signed connection rate. Colors indicate individuals, and triangles indicate family members. The signed connection rate was evaluated no more than two seconds after stimulus onset, which was evaluated whenever the neuron with the largest overall spike count fired. **[g]** Rate-coded manifold projections comparing the same recording session restricted to four subpopulations of identity-specific predictive neurons. Subpopulations are shown (from left to right): all identity-specific predictive neurons, all identity neurons, all cross-modal invariant “concept cells”, and all MvMM neurons. Colors indicate individual identities listed in legends.

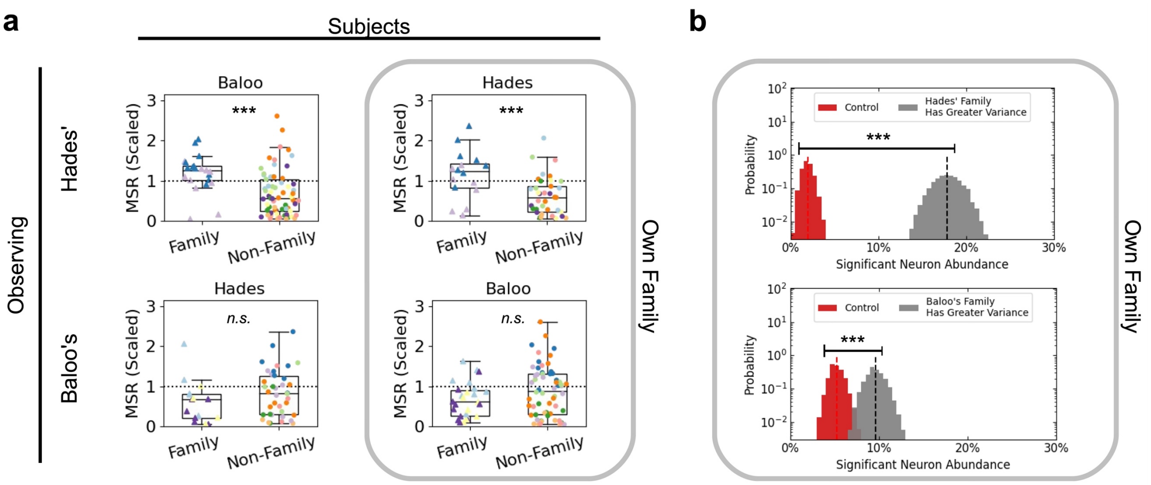


**Figure S12** Significantly different values when subjects were observing the family of (top) Hades and (bottom) Baloo. **[a]** Shown are boxplots of MSR of subjects observing families of other subjects. Significance was computed according to a one-sided Student's t-test consistent with the other subjects viewing the same family, resulting in (top left, $N_{identities} \geq 20$) $p<0.001$, (top right, $N_{identities} \geq 16$) $p<0.001$, (bottom left, $N_{identities} \geq 14$) $p=0.102$, and (bottom right, $N_{identities} \geq 26$) $p=0.055$. Gray box indicates subjects were observing their own families. **[b]** Histograms showing the relative abundance of neurons with significantly larger variance of signed connection rate for the subject's own family relative to other conspecifics according to Fligner-Kileen's test ($p<0.01$). Variance of signed connection rate was computed from the reference neuron to each neuron. Control was a random shuffle of the labels for each neuron. Distributions were determined via bootstrap. Dotted lines indicate the mean values for Hades viewing her own family (left, 18±3% out of N=610) and Baloo viewing her own family (right, 10±2% out of N=822), which both exhibited significantly more significant neurons than control (left, 2.0±1.1% out of N=610; right, 5.2±1.5% out of N=822) according to Student's t-test ($p<0.001$, $N_{bootstrap}=10{,}000$). Uncertainty indicates 95% confidence intervals of the mean. Gray box indicates subjects were observing their own families. Bin width is 0.5%.

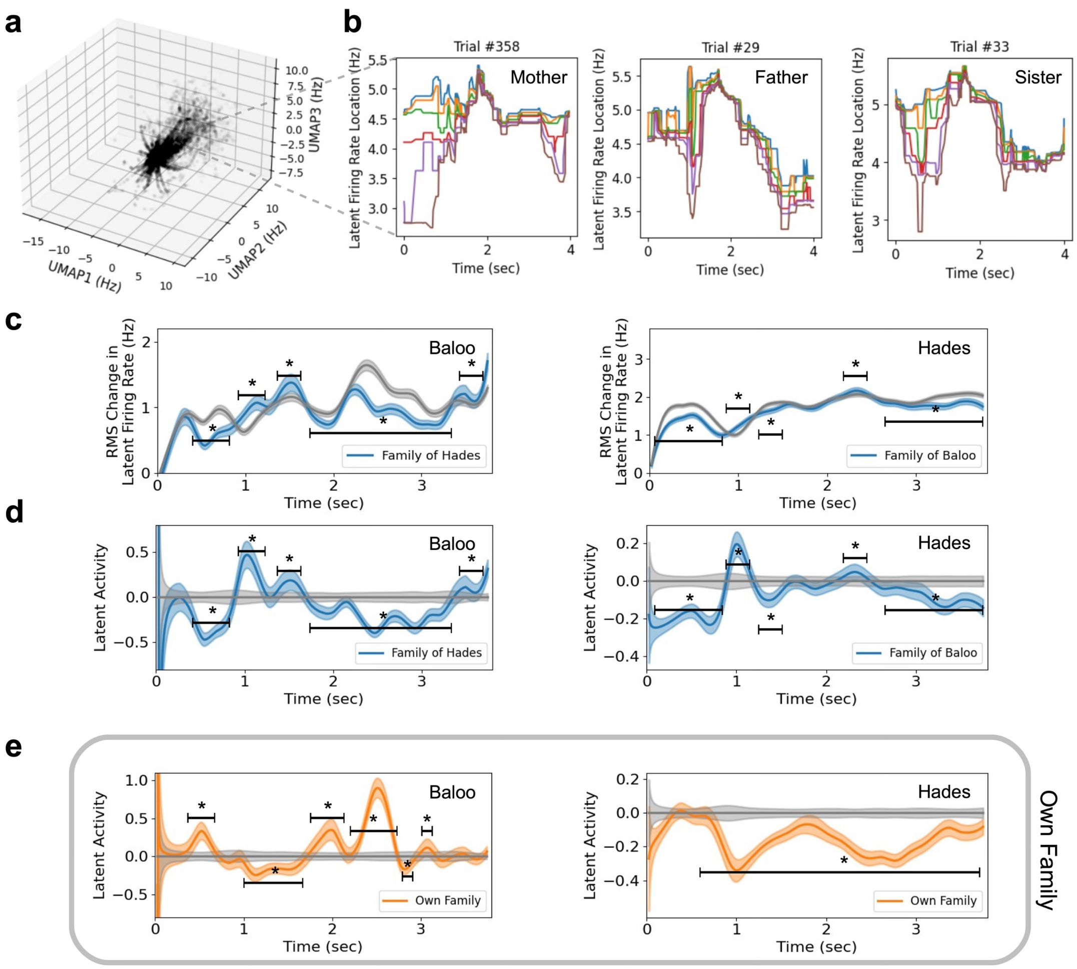


**Figure S13 Quantification of latent activity [a]** Shown are the first three axes of the six dimensional latent firing rate, which was an unsupervised manifold projection of the absolute value of the signed connection rate from the same neuron with the largest overall spike count (i.e. the reference neuron) to all neurons that appeared approximately symmetric (defined in Methods). **[b]** Shown are time traces of our latent firing rate for an exemplary trial from each of three family members of Baloo. Each color represents one dimension. The order of dimensions is consistent between panels. **[c]** Root mean squared (RMS) change in latent firing rate *versus* time averaged over all recording sessions from subjects (left) Baloo and (right) Hades. Traces average over identity-match trials showing (blue) the family members of the subject and (gray) all conspecifics. **[d]** Latent activity *versus* time for (left) Baloo and (right) Hades. Latent activity traces were computed as the ratio of the RMS change in latent firing rate to control minus one. Control was RMS change in latent firing rate averaged over all identity-match trials. **[e]** Latent activity *versus* time for (left) Baloo and (right) Hades viewing their own family. Control was as in [d].

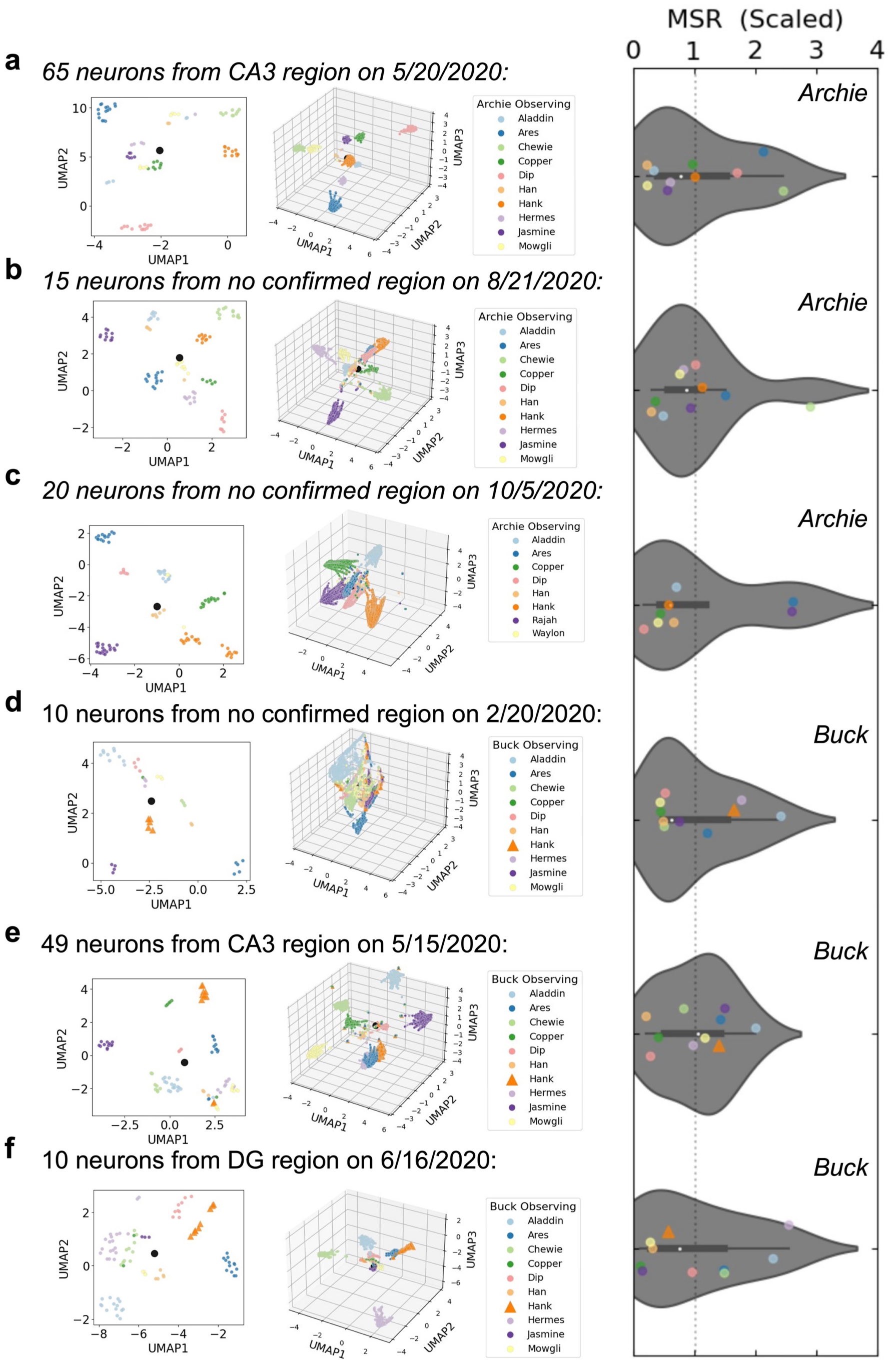


**Figure S14 [a-f]** Manifold projections comparing three different recording sessions conducted on different observers, Archie [a-c] and Buck [d-f]. Shown are rate-coded projections (left), event-coded projections (middle), and MSR computed from the event-coded projections (right). Colors indicate individual identities listed in legends.

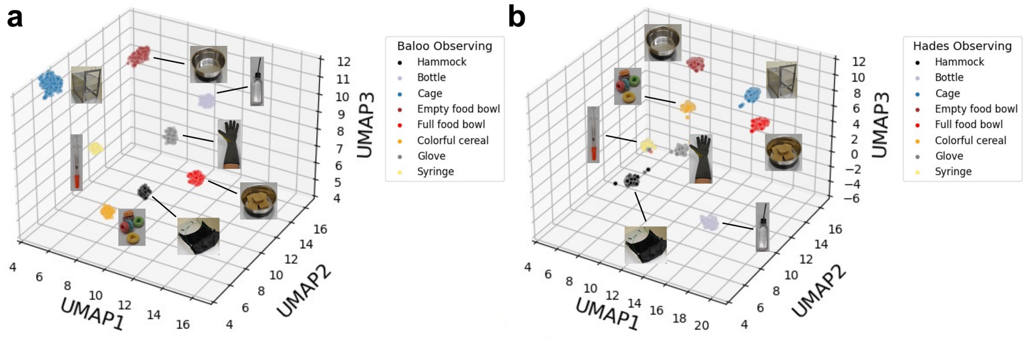


**Figure S15 Separability of socially-agnostic categories** [a-b] Event-coded representations of inanimate objects from the laboratory setting. Separation of socially-agnostic categories are shown in marmoset hippocampus for two subjects, [a] Baloo and [b] Hades. Visual images from each of these object categories was presented to subjects using the same stimulus presentation protocol as for the unimodal stimuli while recording single neuron activity in marmoset hippocampus from two marmosets. Likewise, we performed the same signed-connection rate analysis and input those data into UMAP using the same data analysis pipeline as described for analyses presented in Figure 4.

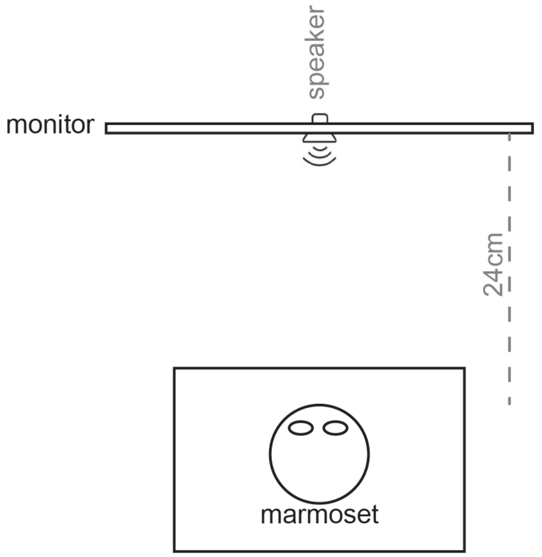


**Figure S16. Schematic drawing of experimental setup.** In an anechoic chamber, marmoset subjects were seated, positioned 24 centimeters away from a monitor and a speaker. The speaker was located just below the monitor.

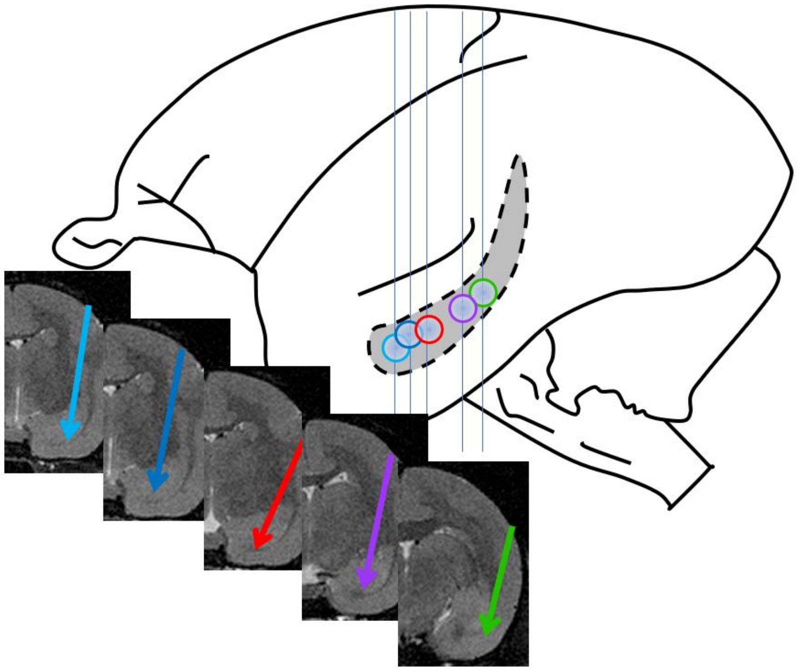

**Figure S17. Anatomical locations of microwire bundles across animals.** Arrows on MRI cross-sections indicate trajectory of each microwire brush array in marmoset hippocampus. Each color indicates a different animal's array. Circles correspond to anterior-posterior position.
